# ^UFM^Track: Under-Flow Migration Tracker enabling analysis of the entire multi-step immune cell extravasation cascade across the blood-brain barrier in microfluidic devices

**DOI:** 10.1101/2023.01.04.522827

**Authors:** Mykhailo Vladymyrov, Luca Marchetti, Sidar Aydin, Sasha Soldati, Adrien Mossu, Arindam Pal, Laurent Gueissaz, Akitaka Ariga, Britta Engelhardt

## Abstract

The endothelial blood-brain barrier (BBB) strictly controls immune cell trafficking into the central nervous system (CNS). In neuroinflammatory diseases such as multiple sclerosis, this tight control is, however, disturbed, leading to immune cell infiltration into the CNS. The development of in vitro models of the BBB combined with microfluidic devices has advanced our understanding of the cellular and molecular mechanisms mediating the multi-step T-cell extravasation across the BBB. A major bottleneck of these in vitro studies is the absence of a robust and automated pipeline suitable for analyzing and quantifying the sequential interaction steps of different immune cell subsets with the BBB under physiological flow in vitro.

Here we present the Under-Flow Migration Tracker (^UFM^Track) framework for studying immune cell interactions with endothelial monolayers under physiological flow. We then showcase a pipeline built based on it to study the entire multi-step extravasation cascade of immune cells across brain microvascular endothelial cells under physiological flow in vitro. ^UFM^Track achieves 90% track reconstruction efficiency and allows for scaling due to the reduction of the analysis cost and by eliminating experimenter bias. This allowed for an in-depth analysis of all behavioral regimes involved in the multi-step immune cell extravasation cascade. The study summarizes how ^UFM^Track can be employed to delineate the interactions of CD4^+^ and CD8^+^ T cells with the BBB under physiological flow. We also demonstrate its applicability to the other BBB models, showcasing broader applicability of the developed framework to a range of immune cell-endothelial monolayer interaction studies. The ^UFM^Track framework along with the generated datasets is publicly available in the corresponding repositories.

**Author summary:** Immune cells continuously travel through our body to perform immune surveillance. They travel within blood vessels at a very high speed and slow down upon reaching their target organ by the sequential interaction with different adhesion and signaling molecules on the vascular endothelial cells.

The study of molecular mechanisms mediating this multi-step extravasation of immune cells has been significantly advanced by in vitro cultures of microvascular endothelial cell monolayers. The dynamic interaction of the immune cells with endothelial monolayers can be imaged over time in vitro in microfluidic devices under physiological flow. The 2-dimensional structure of the endothelial monolayer allows for reliable visualization of the extravasation process required for the study of the molecular mechanisms involved. The manual analysis of the acquired imaging data is time-consuming and prone to experimenter error. Analysis automation is, however, hampered by the similar appearance of the unlabeled immune and endothelial cells and by the flow causing rapid immune cell displacement.

Here we introduce ^UFM^Track, the under-flow migration tracker framework allowing for automated analysis of immune cell interactions with microvascular endothelial cells under flow in vitro. ^UFM^Track performs comparably to the manual analysis of an experienced researcher, eliminates experimenter’s bias, and improves the accuracy of the immune cell tracking. Taken together, ^UFM^Track sets the stage for scalability of in vitro live cell imaging studies of immune cell extravasation.

## Introduction

Immune cells continuously travel throughout our body as a means of immune surveillance. Moving within the bloodstream allows for their fast transport to even distant sites but requires extravasation once they have reached their target organ. Immune cell extravasation across the vascular wall is a multi-step process regulated by the sequential interaction of different signaling and adhesion molecules on the endothelium and the immune cells (1,2). These molecular interactions mediate distinct sequential steps, namely tethering and rolling to reduce travel speed, shear- resistant arrest, polarization and crawling of the immune cell on the luminal surface of the endothelium, and finally, immune cell diapedesis across the endothelial layer (2–4).

The precise molecular mechanisms mediating the multi-step immune cell extravasation in each organ depend on the immune cell subset but also the specific characteristics of the vascular bed. For example, in the central nervous system, the endothelial blood-brain barrier (BBB) establishes a tight barrier that strictly controls the transport of molecules across the BBB, ensuring tissue homeostasis required for neuronal function (2,5). The BBB similarly controls immune cell trafficking into the CNS. Thus, accounting for these special barrier properties, unique characteristics of the multi-step T-cell migration across the BBB have been described. For instance, T cells crawl for very long distances against the direction of blood flow on the surface of the BBB endothelium in search of permissive locations for diapedesis (6–8). Research on T-cell interaction with the BBB has already been successfully translated into therapies in the clinic (9,10).

Exploring the entire multi-step extravasation of immune cells across the BBB has been significantly advanced by making use of in vitro BBB models maintaining their barrier properties, and placing them into microfluidic devices (11). Combined with microscopic setups that allow for in vitro live-cell imaging (Figure 1A-B) of the immune cell interaction with the brain endothelial monolayer under physiological flow over time, the molecular mechanisms mediating the sequential interaction of T cells during extravasation across the BBB have been delineated (2). The live cell imaging is largely performed in the phase-contrast imaging modality (Figure 1C), which does not require establishing fluorescent labels of the imaged cells and avoids potential photo-toxicity of the fluorescent imaging modality. These studies have shown that upon their arrest, T cells polarize and either crawl at speeds between 3 to 10 µm/minute over the brain endothelial monolayer or probe the endothelial monolayer by remaining rather stationary and sending cellular protrusions into the endothelial monolayer (Figure 1D) (12,13). Both behaviors can lead to diapedesis of the T cells across the brain endothelial monolayers. This process lasts at least 3 to 5 minutes, with some immune cells observed to protrude and retract several times prior to finalizing a prolonged diapedesis process to the abluminal side of the brain endothelial monolayer (8,14). Finally, T cells that have successfully migrated across the brain endothelial monolayer usually continue to migrate underneath the endothelial monolayer (15,16).

**Figure 1.**
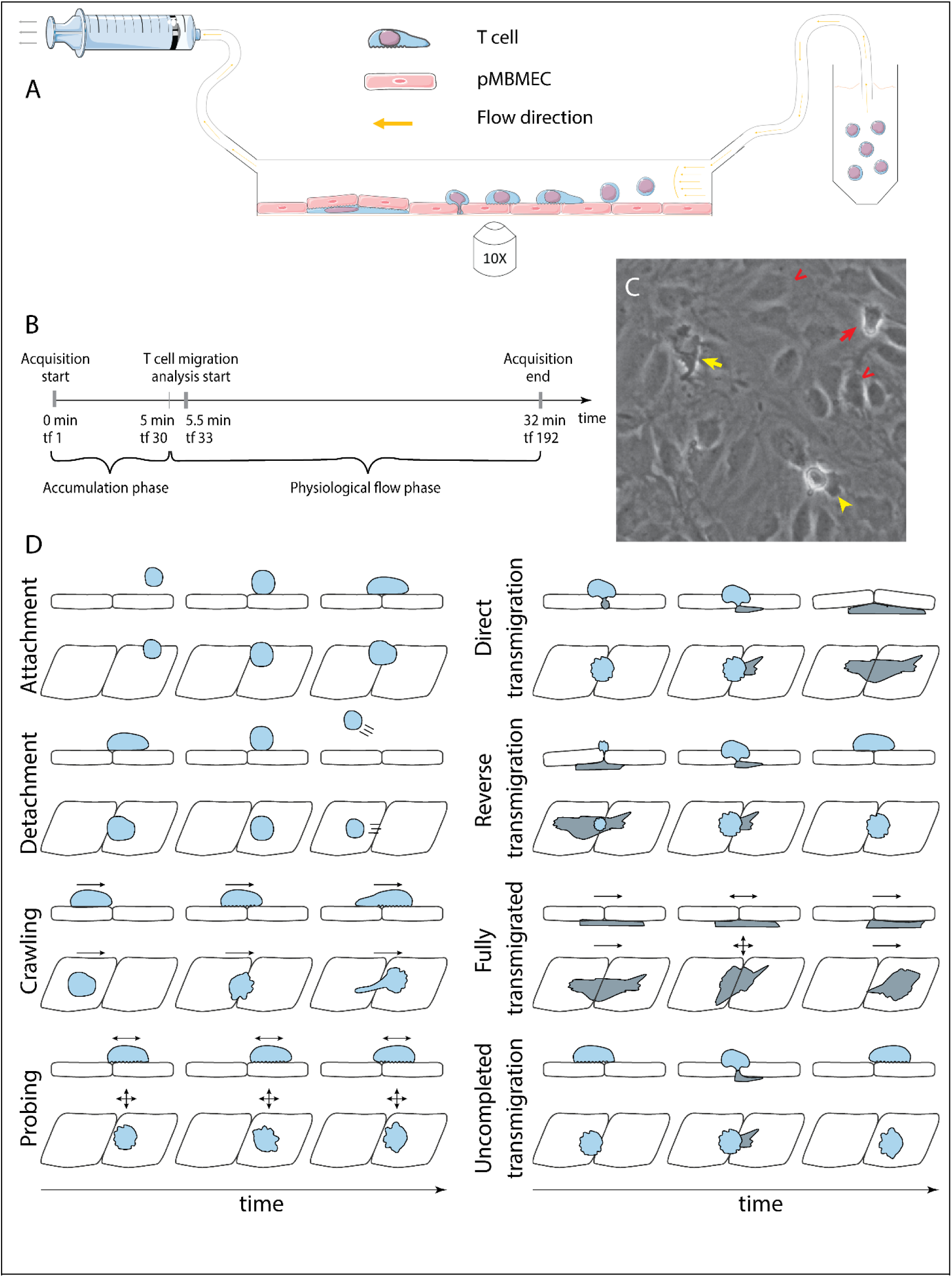
In vitro analysis of the multistep cascade of T-cell migration across the BBB model under physiological flow. A. In vitro under-flow assay setup. T cells were perfused on top of the pMBMEC monolayer, and their migration under flow was observed using phase-contrast imaging modality. Imaging was performed with a time step of 10 seconds/timeframe. B. In vitro flow assay timeline. During the accumulation phase under flow with the shear stress of 0.1 dynes/cm2, T cells adhered to the pMBMEC monolayer. After 5 minutes (timeframe 30), the shear stress was increased to 1.5 dynes/cm2, leading to rapid detachment of not firmly adhering T cells. Analysis of the post-arrest T-cell behavior was thus starting at 5.5 minutes (timeframe 33). tf = timeframe. C. Example of phase-contrast imaging data. Red arrow – crawling T cell; yellow arrow – fully transmigrated T cell; yellow arrow-head – transmigrated part of a partially transmigrated T cell; red V arrowheads – pMBMECs. D. Schematic representation of distinct T-cell behavior regimes detected and analyzed using the developed UFMTrack framework. Crawling cells migrate continuously while probing cells interact with the pMBMECs and move around the interaction point within two cell-size (20 *μm*) as indicated by the arrows. Side and top views are shown.

The data analysis of these “in vitro flow assays” requires time-consuming offline frame-by-frame analysis of the imaging data by individual experimenters, in which the dynamic interactions of each individual immune cell has to be followed over the entire time of the assay manually and assigned to specific categories. Such analysis is tedious, and accurately assigning the different T-cell behaviors requires experience. Thus, this manual analysis is prone to inevitable errors and subjective judgments of the various experiments. Furthermore, the time-consuming manual T-cell tracking limits the number of events that can be studied and, thus, the statistical power of the analysis.

Automation of the analysis of the recorded multi-step T-cell extravasation across the BBB in the microfluidic device would thus be highly desirable. It is, however, hampered as these assays are usually performed with unlabeled cells and imaged by phase contrast (Figure 1C), which poses a challenge due to the similar grayscales and morphology of the immune cells interacting with the brain endothelial cells. Further challenges include perfused on top T cells that, in the presence of shear flow, instantly appear within the field of view (FoV) and are either suddenly displaced over a certain distance or completely detached and washed away (Figure 1D). Proper analysis of these events is mandatory for reliable T-cell tracking but also with respect to the study of the overall avidity of the dynamic T-cell interaction steps with the underlying brain endothelium. Thus, it was compulsory to establish a tracking solution that accounts for the effect of the flow on the migrating cells and the distinct migration regimes.

Segmenting of cells imaged in phase contrast migrating on top of cellular monolayer of similar appearance is a challenging task. Furthermore, the analysis of T cell interactions requires detection and quantification of the transmigration. While recent advances in computer vision established several frameworks for detection (17–20) or segmentation (21,22), no tool is readily available to complete both tasks simultaneously.

There is a large number of powerful cell tracking solutions developed that focus on different aspects of cell migration, and are embedded in different frameworks or solutions. The general-purpose Trackpy (23) allows for simple particle tracking and is not suitable to track cells migrating under flow. It is used for the tracking of cells imaged in phase contrast in the Usiigaci framework (24). The widely used TrackMate (25) plugin for ImageJ (26) allows for tracking of particles performing either the Brownian motion or a linear motion, but not the combination of two. Instant appearance or disappearance as well as the resolving tracks of under-segmented cells cannot be easily established in this framework. Similarly, CellTraxx (27) performs matching of cells in the adjacent frames, prohibiting it’s use for tracking of under-segmented, intersecting cells or under flow. The Bayesian Tracker (btrack) framework (28) is using spatial information as well as appearance information for track linking focusing on the cell divisions required for cell lineage tracing.

Here we introduce the developed under-flow migration tracker (^UFM^Track) framework that systematically addresses the hurdles mentioned above, allowing it to perform automated tracking and analysis of cell-cell interactions imaged in phase contrast. We also show a successful implementation of ^UFM^Track to build an analysis pipeline for T-cell interactions with brain microvascular endothelial cells in vitro under physiological flow. ^UFM^Track reaches 90% T-cell tracking efficiency, performing comparably to manual analysis while eliminating the experimenter’s bias and improving the accuracy of T-cell tracking. Therefore, it enables significant savings in the labor force and time for data analysis.

## Results

To design and develop the automated T-cell under-flow migration analysis framework ^UFM^Track framework presented here, we made use of in vitro imaging datasets following T-cell migration across primary mouse brain microvascular endothelial cells (pMBMECs) under physiological flow in vitro. The framework combines three components: T-cell segmentation and transmigration detection, T-cell tracking under flow, and analysis of each of the multistep T-cell migration cascade steps. Segmentation and transmigration detection of the T cells, migrating on the pMBMECs, is performed with a 2D+T U-Net-like convolutional neural network (21). The T-cell under-flow tracking algorithm was formulated as a constrained optimization problem.

Next, we describe the methods and algorithms employed to develop ^UFM^Track. Links to the code and the datasets used for model training can be found in the Data Availability section.

### I. T-cell segmentation

Reliable cell segmentation is crucial for reliable cell tracking. In the phase-contrast imaging modality, it was impossible to achieve reliable differentiation between T cells and endothelial cells of the pMBMEC monolayer based on pixel intensity. For detection of T-cell transmigration across pMBMEC monolayers (diapedesis), sufficiently reliable cell segmentation of the transmigrated T cells was also required. To achieve this, we have designed 2D and 2D+T U-Net-like (21) fully convolutional neural network-based models for multitask learning. The models were trained to predict three maps: cell probability, the probability that the T cell is below the pMBMEC monolayer, and cell centroids. The models performed predictions based on the grayscale of the respective phase- contrast images in the case of the 2D model or sequences of 5 timeframes in the case of the 2D+T model. The models were implemented in TensorFlow (29). The training was performed on a dataset corresponding to 154 Megapixels of raw image data using the annotation mask for T cells (“T cell mask”) and the mask of the transmigrated part of the T cells (“transmigration mask”), the centroids map, and the weight map (Figure 2). For validation 37 Megapixels of raw imaging data were used.

**Figure 2.**
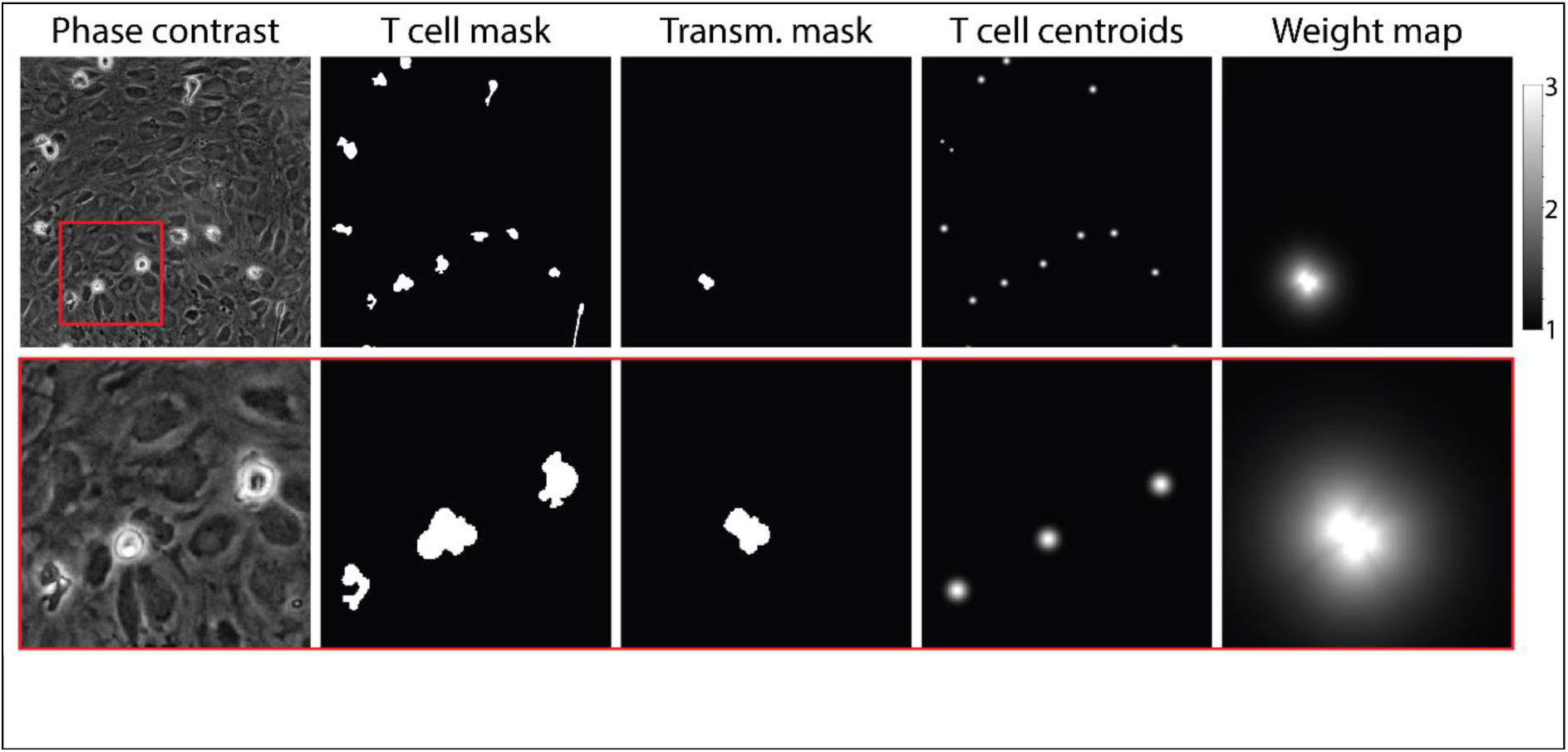
Example of data used for training of the T-cell segmentation model. From left to right: Phase-contrast microscopy image of T cells migrating on top of the pMBMEC monolayer; annotated T cell mask; annotated mask of transmigrated part of T cells; T cell centroids; *W*_*cell*_ weight map. The bottom row shows zoom-in on the highlighted area.

The models have approximately 3M and 8M parameters correspondingly, and the model architectures are summarized in Supplementary Tables 1, 2. Details on models’ implementation, training, and image processing can be found in the “Segmentation models” section of the Supplementary Material.

In the T cell mask prediction task, the 2D+T model outperformed the 2D model by a notable 9% according to the average precision (AP) metric (Table 1). At the same time, for the transmigration mask, which is much more difficult to infer, the 2D model performance reached only 54%, which is insufficient for reliable detection of T cells migrating across the pMBMEC monolayer. In this task, the 2D+T model outperformed the 2D model by 32% AP. We observed that the 2D+T model was sensitive to frame misalignment, leading to false positive detection of transmigrated T cells. Thus, to generate the preliminary T-cell masks used for frame alignment and histogram normalization, we employed the 2D model. Afterward, for the T-cell segmentation and transmigration detection, we employed the 2D+T model, followed by a watershed algorithm based on the inferred T-cell mask and the cell centroids as seed points. A stitched phase-contrast image sequence can be seen in Supplementary Video 1 and overlayed with the segmented cells and highlighted transmigration mask in Supplementary Video 2.

**Table 1.**
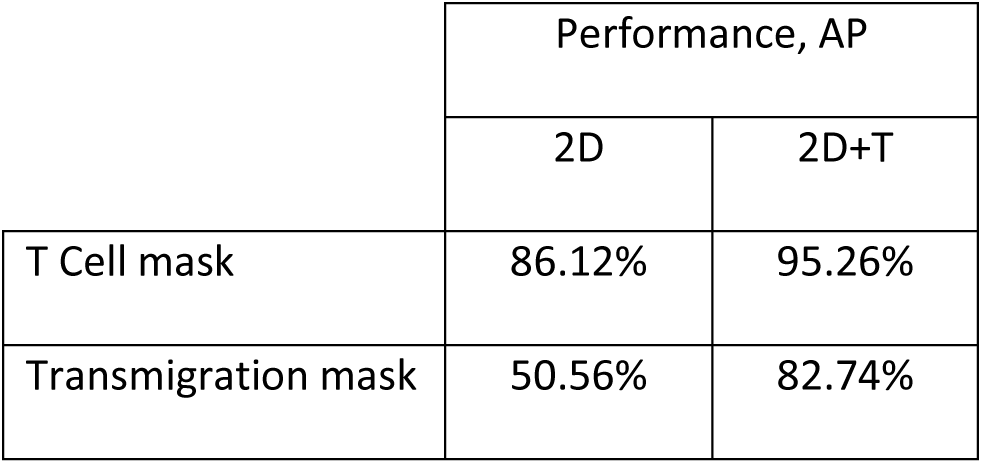
Comparison of the performance of the 2D and 2D+T-cell segmentation models.

After segmentation, we suppressed the noise by discarding small objects with an area below 30 pixels (the mean area of a T cell is about 400 pixels). For each T cell, we evaluated its coordinates as geometric mean coordinate, angle of the longer axis, and elongation 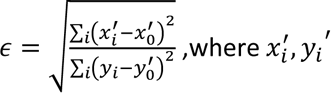 are projections of the *i*-th T cell mask pixel coordinates on the orthogonal long and short axes of the T cell, and *x*^′^_0_, *y*^′^_0_ are the corresponding projections of the geometrical center of the T cell. The summation is performed over the area covered by the mask of the corresponding T cell. Additionally, we evaluated the mean T cell probability 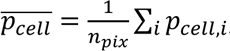, mean transmigration probability 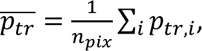, and cell transmigration coefficient as 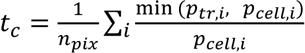 number if cell pixels and *p*_*cell*,*i*_, *p*_*tr,i*_ are the T cell and transmigration probabilities for *i*-th pixel in the predicted T cell mask.

### II. T-cell tracking

The presence of flow causes several types of discontinuous events that were managed with a specialized tracking algorithm. These are: 1) the sudden appearance of T cells in the field of view (FoV), 2) the sudden detachment of a T cell followed by its disappearance from FoV, and 3) the displacement over significant distances (up to hundreds of micrometers) of T cells that do not adhere firmly to the pMBMECs. The primary inspiration for our approach was the Conservation Tracking algorithm (30). We could consistently reconstruct the T-cell tracks by performing global optimization constrained to controlled probabilities of T-cell appearance, disappearance, and displacement due to the flow at every time point. The standard tracking algorithms like tracking by association is either unable to detect track links with high displacement and thus reconstructs several disconnected segments of a cell track, or leads to a high number of false track links. Our procedure favors the reconstruction of long T-cell tracks while at the same time allowing for tracking of the T cells which detach or are displaced by the flow (“accelerated T-cell movement”) over a longer distance (> 8 *μm*) with speed significantly higher than their crawling speed.

The tracking consists of four main steps: linking, searching for track candidates, global track consistency resolving, and resolving the track intersections. The inputs for the tracking are the segmented cells and the cell centroids (Figure 3A). The centroids and the cell proximity are used to quantify under-segmentation by introducing the node multiplicity (Figure 3B, see section T-cell tracking of the supplementary information for details). At the linking step, we identified all possible connections of T cells between the timeframes (Figure 3C).

**Figure 3.**
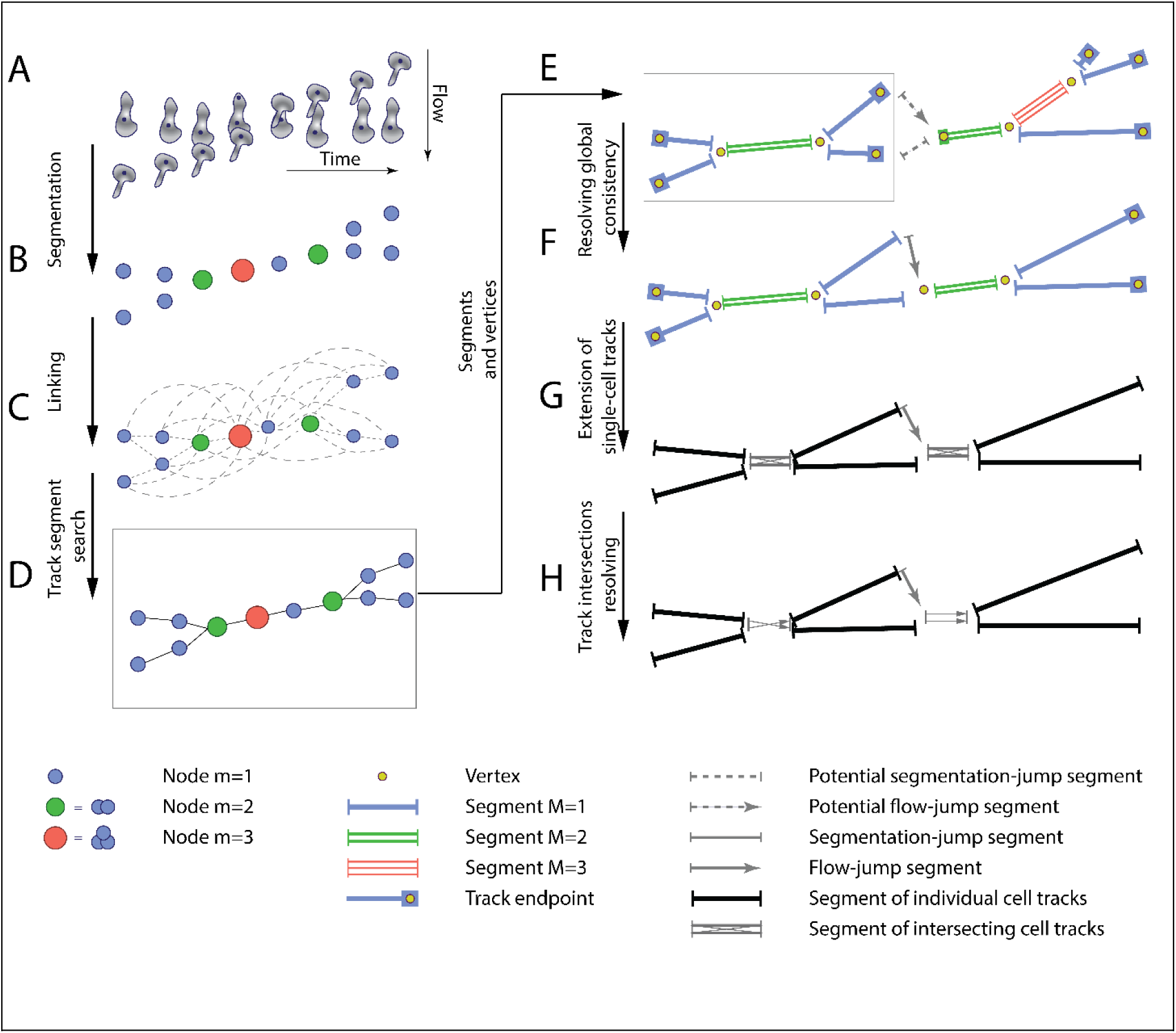
The T-cell tracking pipeline. A. Segmentation and centroids of two migrating T cells under flow over time. B. Nodes corresponding to segmented T cells. Node sizes and colors denote the multiplicity estimate of a node. C. Nodes, together with links between adjacent in space and time nodes, form a graph. D. Selected links are obtained with global optimization on the graph. E. Graph of segments and vertices obtained according to node multiplicity. F. Resolving global segment multiplicity consistency with global optimization on the graph. Additionally, a search for rapid displacement of segmented T cells due to under-segmentation or flow is performed. G. Extension of tracks of individual T cells into the track intersection. H. Resolving track intersections.

Next, during the search for track candidates, we aimed to find continuous track segments of T cells or under-segmented groups of T cells crawling on top of or below the pMBMEC monolayer without accelerated movement segments on the T cell track characterized by rapid T cell displacements. The whole dataset of T cells across all time points was represented as a graph. Each vertex corresponds to a T cell (multiplicity m=1) or a group of potentially under-segmented T cells (m>1). The vertices are connected according to links obtained at the linking step. Track segments were found by performing global optimization to find consistent connectivity of vertices across the time points (Figure 3D) by employing an approach inspired by the conservation tracking algorithm summarized in Schiegg et al. (30). Optimization was performed using the CP-SAT constrained optimization procedure using the open source OR-Tools library (31).

Next, resolving global track consistency was performed. In this step, the scope of T cell tracking is shifted from individual nodes (representing T cells at particular timeframes as well as groups of under-segmented T cells) to the track segments (unambiguous sequences of nodes) and vertices at the endpoints of the segments. These are track start and end points, points of merging and separation of track segments in case of under-segmentation, as well as ambiguous points on a track. The latter was identified by sudden T cell displacement, a hallmark of detaching and reattaching T cells and T cells transitioning from properly segmented to under-segmented T cells or vice versa (Figure 3E). This was followed by the search for potential missing track segments due to accelerated T cell movement under flow and missing links in the track crossing points. Both are characterized by significant displacement lengths such that they were not detected during the linking step. Thus we will refer to both as “jumps” in this section. We have also eliminated short and therefore unreliable segments, as well as segments whose multiplicity was found to be *M* = 0 (Figure 3F). Afterward, we separated the segments into two categories, namely segments with multiplicity *M* = 1, i.e., tracks of isolated T cells, and segments with multiplicity *M* > 1, i.e., tracks of under-segmented groups of T cells, where tracks of several T cells intersected. Finally, we extended the tracks of isolated T cells into the intersection if the branching vertex had a multiplicity of *n* and was splitting in exactly *n* segments with multiplicity *M* = 1 (Figure 3G).

Lastly, we resolved the track intersections to obtain reliable T cell tracks, i.e., identifying track segments corresponding to the same T cell before and after the under-segmented track region (Figure 3H). Detailed information on the developed under-flow T cell tracking algorithm is given in the “T-cell tracking” section of the Supplementary Material.

### III. T-cell migration analysis

Next, we performed the T-cell migration analysis based on the reconstructed T cell tracks. We selected tracks inside the fiducial area of the FoV, namely coordinates of the T cell at all timepoints along the track were located at least 25 µm away from the bounding box enclosing all segmented T cells. Next, tracks of T cells touching another T cell at the end of the assay acquisition were excluded since T cells directly adjacent to each other can hide the start of T cell transmigration across the pMBMEC monolayer and thus compromise correct detection and quantification of this step.

Additionally, only tracks with T cells were assigned in at least 6 timeframes during the physiological flow phase. We also require T cells to be assigned for at least 75% of timeframes along the track.

Under-segmented parts of T cell tracks were not considered. Examples of selected tracks can be seen in Supplementary Video 3.

After exclusion of the tracks of the cellular debris, which were misclassified as T cells during the segmentation step, using a dedicated “not-a-T-cell” classifier, we performed identification of transmigration (Supplementary Figure 5), probing and crawling migration regimes, as well as accelerated movement along the T cell track. Detailed information on the analysis of the T cell tracks is given in the “T-cell migration analysis” section of the Supplementary Material.

Finally, for each track, we evaluated motility parameters for each of the following migration regimes: probing before the transmigration, crawling before the transmigration, all crawling above pMBMECs monolayer including T cell crawling segments after the first transmigration attempts, all crawling below pMBMECs monolayer, whole T-cell track excluding accelerated movement and tracking inefficiency regions, as well as whole T-cell track. Specifically, we evaluated the following T-cell motility parameters: duration of each migration regime, the total vector and absolute displacements, the migration path length, the average migration speed (displacement over time), average crawling speed (path length over time), and finally the mean and standard deviation of the instantaneous speed. We evaluated migration time, displacement, and average speed for the accelerated movement regime.

### IV. Analysis of trafficking datasets

Having developed a full pipeline based on the ^UFM^Track framework for automated analysis of the multi-step T-cell migration cascade across the BBB model, we next aimed to compare the results of the automated analysis with previous studies, assess the capacity of the framework to gain novel insight into T-cell migration under flow, and evaluate its performance as compared to manual analysis.

#### 1. CD4^+^ T cell analysis

To this end, we first analyzed a total of 18 imaging datasets each corresponding to 32 minutes recording dedicated to understanding the multi-step migration of CD4^+^ T cells across non-stimulated (n= 6), TNF stimulated (n=5), and IL1-β-stimulated (n=7) pMBMEC monolayers under physiological flow with the developed pipeline.

In these in vitro live cell imaging datasets, phase-contrast imaging was performed as described in the “Data acquisition” subsection in the Materials and Methods section. The T-cell accumulation phase corresponding to the shear stress of 0.1 dynes/cm^2^ lasted for 32 timeframes (5 minutes), followed by conditions of physiological flow (shear stress 1.5 dynes/cm^2^) for the subsequent 160 timeframes (27 minutes). The shear stress is controlled by an automated syringe pump. After increasing to physiological flow rates (12), a significant number of T cells detached from the pMBMEC monolayers, with higher numbers of T cells detaching from non-stimulated pMBMEC monolayers when compared to those stimulated with pro-inflammatory cytokines as detected by the distribution of the track ending times between 5 and 30 minutes. (Figure 4A).

**Figure 4.**
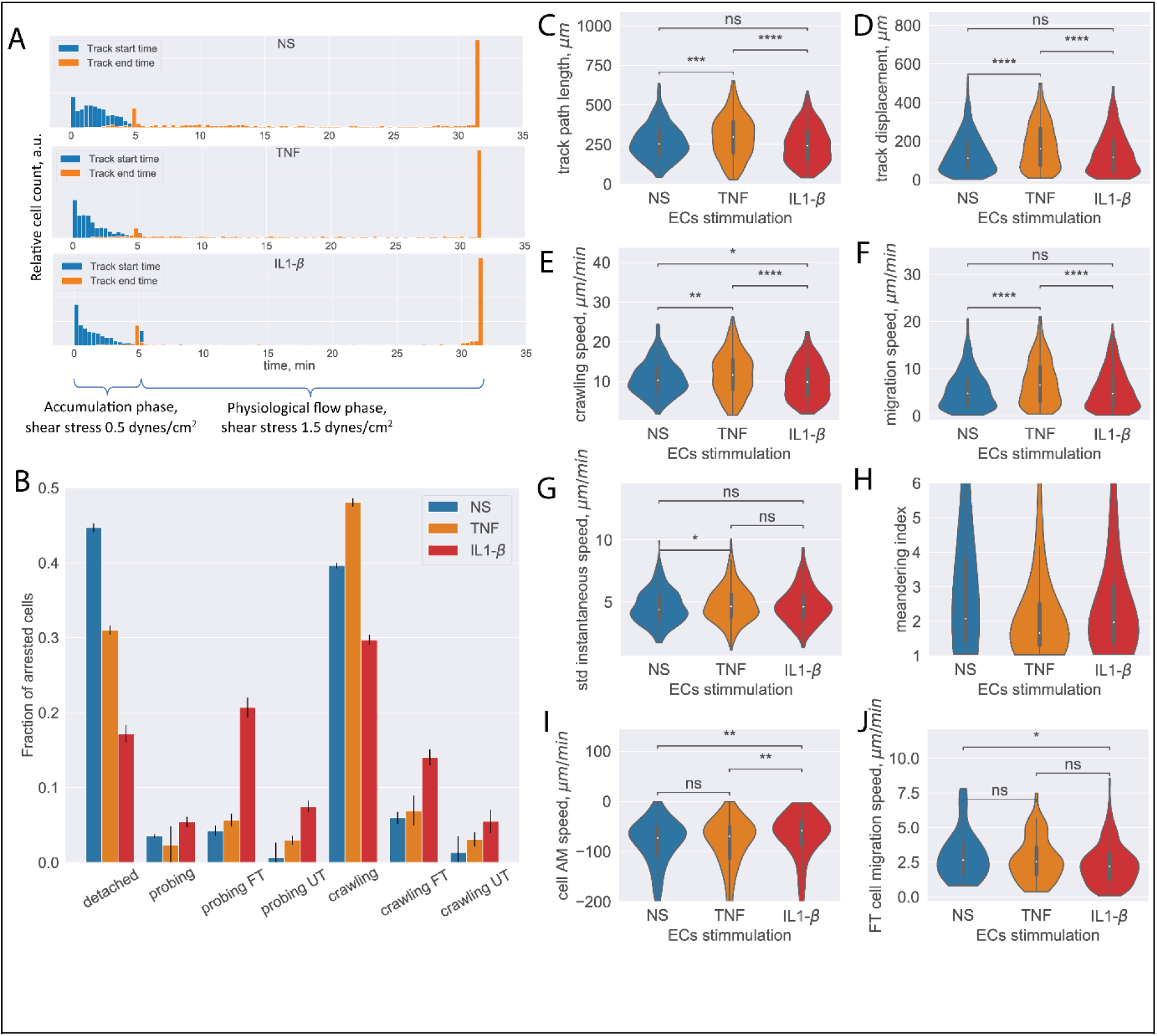
Analysis of CD4^+^ T-cell tracks. A. Start and end time distribution of CD4^+^ T-cell tracks on non-stimulated or TNF or IL-1β stimulated pMBMECs. ^UFM^Track correctly captures increased T-cell detachment from the non-stimulated pMBMECs after 5 minutes when the flow is increased to physiological levels. B. Quantification of CD4^+^ T-cell behavior in the respective categories obtained on non-stimulated and TNF and IL-1β stimulated pMBMECs. Error bars show the statistical error of the mean. (see text for details). C-H. Motility parameters of the crawling CD4^+^ T cells were obtained for the three endothelial stimulatory conditions. Distributions of T-cell path length (C), displacement (D), crawling speed (path/time) (E), migration speed (displacement/time) (F), variability of instantaneous T-cell crawling speed along the track (standard deviation, G), and meandering index (H). I. Distribution of CD4^+^ T-cell accelerated movement (AM) speed is a proxy metric for the T-cell adhesion to the healthy or inflamed endothelium. J. Migration speed distribution of the transmigrated CD4+ T cells. Stimulation applied to the luminal side of pMBMECs affects T-cell migration at the abluminal side of pMBMECs after their transmigration. FT - T cells performed full transmigration; UT – T cells performed uncompleted transmigration.

Analyzing these datasets with the established ^UFM^Track framework, the T-cell behavior type for each detected T-cell track and the aggregated T-cell behavior statistic were obtained for the stimulated and non-stimulated pMBMEC monolayers (Figure 4B). The total number of analyzed T cell tracks is n=971, n=780, and n=1218 for the non-stimulated, TNF stimulated, and IL1-β stimulated pMBMEC monolayers correspondingly. The data obtained with ^UFM^Track were in accordance with our previous observations obtained by manual frame-by-frame analysis (12,14,32). We further obtained the data for T-cell migration speed, T-cell displacement, and T-cell path lengths for the crawling T cells, as these are the primary cell motility parameters. As shown in Figure 4C-H, we observed statistically significant differences in the mean T-cell motility parameters depending on the pMBMECs stimulation condition. Furthermore, we observed differences in the shape of the distribution of T-cell crawling speeds, consistent with our previous reports of underlying differences in the mechanisms mediating T-cell crawling on non-stimulated versus cytokine-stimulated pMBMECs (32). While not statistically significant for the IL1-b and with a low significance for the TNF stimulations, an increased variance of the T cell instantaneous speed on stimulated compared to non-stimulated pMBMECs is observed(Figure 4G). This suggests that T-cell crawling is often interrupted by T-cell recognizing specific cues on the stimulated endothelium. The T cell meandering index (MI) distribution was high on non-stimulated and lower on stimulated pMBMECs (Figure 4H), underscoring that cytokine stimulation enhances directed T-cell movement on the pMBMEC monolayer.

Since the analysis was performed automatically with ^UFM^Track, it allowed for deeper insights into the T-cell migration behavior on the pMBMEC monolayers than obtained by manual frame-by-frame analysis. Specifically, the detection of accelerated movement by ^UFM^Track enables the researcher to quantify the kinetics of individual T-cell detachment from the endothelium rather than simply quantifying a bulk T-cell detachment rate. This detachment kinetics is reflected in the average speed experienced during accelerated movement, which is significantly lower for the IL1-b condition (Figure 4I), suggesting the need to break more bonds with the endothelium when pro-inflammatory cytokines are present.

As our ^UFM^Track workflow achieves sufficient segmentation efficiency to detect T cells below the pMBMEC monolayer and T-cell transmigration across the pMBMEC monolayer, it also enables investigation of T-cell movement after the transmigration step. In Figure 4J, we show the distribution of the migration speed for the transmigrated CD4^+^ T cells. Interestingly, cytokine stimulation of pMBMECs, although applied from the luminal side, also affected T-cell movement at the abluminal side of the pMBMEC monolayer. While in the microfluidic device used in the present study, the migration of T cells below the pMBMEC monolayer may not be biologically relevant, this analysis option will be valuable for future studies involving multilayer in vitro BBB models that include the vascular basement membrane in addition to pericytes and astrocytes to mimic the entire neurovascular unit (8,33).

Importantly, our ^UFM^Track automatically detects and characterizes unusual events, such as reverse T-cell transmigration, that are easily overlooked with manual counting. We do not present the statistics here, as only a few such events were observed. However, when applied to the multilayer BBBs in forthcoming in vitro models, systematic detection and analysis of these rare events will be important to understand T-cell migration in immune surveillance.

#### 2. CD8^+^ T cells analysis

As we have previously shown that the multi-step T-cell extravasation across pMBMEC monolayers differs between CD4^+^ and CD8^+^ T cells (13), we next analyzed two datasets studying the multi-step migration of CD8^+^ T cells across non-stimulated (NS) and TNF/ interferon-gamma (TNF/IFN-γ) stimulated pMBMEC monolayers under physiological flow over 161 timeframes (27 minutes). The CD8^+^ T cells were slightly different in size and appearance when compared to the CD4^+^ T cells used for training the segmentation model. To benchmark our established ^UFM^Track pipeline in this more difficult configuration, the datasets were first manually analyzed by four experimenters: one advanced experimenter with four years of experience (AdEx) and three inexperienced experimenters who received comparable 2-hour introduction and training (Ex1-Ex3). The analyses were performed on the subset of the acquisition, 161 timeframes long starting from timeframe 31.

Manual cell analysis and tracking were performed as described in the “Materials and Methods” section separately for each of the 8 tiles of the tiled acquisition. Next, all crawling CD8^+^ T cells that did not perform transmigration were manually tracked for the timespan after the flow was increased to the physiological level using the manual tracking in ImageJ.

First, we compared the CD8^+^ T cell behavior statistics obtained manually and by our automated analysis pipeline (Figure 5A-C). The results obtained by the automated analysis were in full agreement with the manual analysis performed by the experienced experimenter. At the same time, the data highlight the variability of the results from the inexperienced experimenters, confirming the potential for significant experimenter bias.

**Figure 5.**
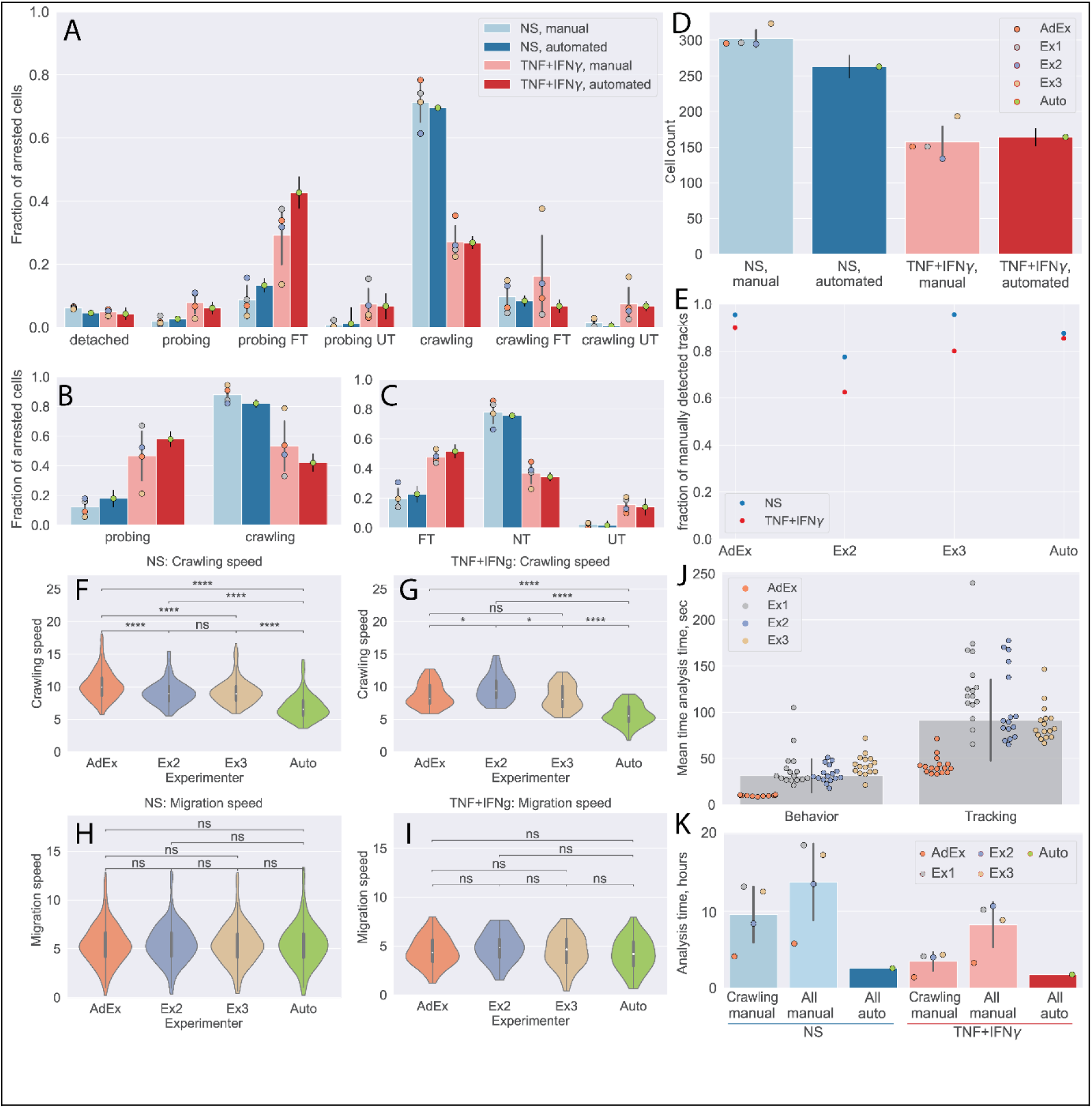
Comparison of automated analysis with ^UFM^Track and manual analysis of the CD8^+^ T-cell tracks. A-C: CD8^+^ T-cell behavior statistic obtained for non-stimulated (NS) and cytokine-stimulated pMBMECs as obtained manually by four experimenters, as well as automatically with ^UFM^Track. A. Quantification of CD8^+^ T-cell behavior in the respective categories obtained on non-stimulated and TNF/IFN-γ stimulated pMBMECs is consistent with results obtained by manual frame-by-frame analysis. Cytokine stimulation of pMBMECs increases T-cell probing behavior (B) and T-cell transmigration rate (C). Error bars show the standard deviation of the manual analysis and the statistical error of the mean for automated analysis. Points correspond to individual experimenters. D. Counts of CD8+ T cells were obtained manually by four experimenters and automatically by UFMTrack. The T-cell detection efficiency is above 90%. Error bars show the standard deviation of the manual analysis and the statistical error of the mean for the automated analysis. Points correspond to individual experimenters. E. Detection efficiency of crawling CD8+ T cell tracks between the manual and automated analysis. F-I. Comparison of CD8+ T-cell crawling speed (path/time) (F, G) and migration speed (displacement/time) (H, I) on non-stimulated (F, H) and cytokine-stimulated (G, I) pMBMECs. The T-cell position assignment error in manual tracking leads to biased crawling speed estimation. J. Comparison of the analysis time (per cell) required for behavior analysis and tracking of CD8+ T cells. K. Total analysis time (per dataset) for behavior analysis and tracking of CD8+ T cells. Comparison is shown for manual analysis with tracking of crawling cells only (Crawling manual), the time estimate for manual analysis with tracking of all cells (All manual), and the in-depth automated analysis of all cell tracks with UFMTrack (All auto). FT - T cells performed full transmigration; UT – T cells performed uncompleted transmigration; NT - T cells did not perform transmigration.

We also show the detection efficiency of T cells by the ^UFM^Track. To achieve this goal, we combined the numbers of CD8^+^ T cells detected in each tile by manual analysis. Due to the overlap between the individual tiles, some CD8^+^ T cells are seen more than once. To obtain an estimate of the total CD8^+^ T cell count detected manually, we scaled the number of CD8^+^ T cells accordingly to the number of detected unique T-cell tracks (see next paragraph). By this approach, we found T-cell detection efficiency to be above 90% (Figure 5D). The increased T-cell density can explain lower efficiency in the NS condition. These findings set the appropriate T-cell density for automated analysis of their migration behavior on pMBMEC monolayers to 200-300 cells per FoV of the size of 3.8 mm^2^ or 50-80 cells/mm^2^.

Next, we compared the T-cells tracks as obtained manually by three experimenters as well as automatically by our ^UFM^Track. To this end, we first matched the tracks in adjacent tiles to obtain the tracks on the whole imaging area, avoiding multiple counts of the same track. This was achieved by pattern matching with initial offsets between tiles obtained by an automatic frame alignment procedure performed as part of the automatic analysis. We considered T-cell tracks observed in different tiles to be tracks of the same T cell if 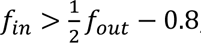, where *f*_*in*_ is the fraction of timepoints along a track at which the distance between the T cells was below 17 µm and *f*_*out*_ was the fraction of timepoints along a track at which distance between cells was above 25 µm. We used the same approach to match the T-cell tracks obtained by each experimenter and the automated ^UFM^Track. To compare performance in an objective manner, we excluded manually obtained tracks that lay outside of the fiducial volume of the automated analysis, as well as T cells that were touching another T cell at the end of the acquisition, as those were also excluded from the automated analysis. We then took as 100% the sum of all T-cell tracks detected manually. We evaluated the fraction of all T-cell tracks detected by each experimenter as well as automatically by our ^UFM^Track.

In Figure 5E, we show that our pipeline achieves a comparable T-cell tracking efficiency when compared to the manual analysis. While its performance was not superior to that of the experimenter with four years of expertise in the analysis of the under-flow datasets, it does perform better than less experienced experimenters.

We also compared the T-cell motility parameters obtained manually and automatically for the non-stimulated and stimulated pMBMECs (Figure 5F-I). Data obtained for the CD8^+^ T-cell migration speed on pMBMECs was comparable between all experimenters and the automated analysis. In contrast, results obtained for the T-cell crawling speed were significantly different between the automated approach and manual analysis. Taking a closer look at the T-cell tracks (Supplementary Figure 6), we readily observed that the manual analysis contains significant errors in the assignment of the T-cell position. This leads to a jittery pattern in the T-cell migration tracks and overestimating the T-cell path and, thus, T-cell crawling speed. Employing our novel automated analysis pipeline eliminated this systematic error in the measurements.

To finally investigate the potential benefit in the time required for automated versus manual data analysis, we examined the time required for a given experimenter to analyze such datasets manually using the Clockify time tracker (34). In Figure 5J, we show the average time investment needed to perform a complete T-cell behavior analysis and T-cell tracking in each imaging tile, as well as on average. Clearly, the experienced experimenter outperforms the inexperienced experimenters by a factor of 3. On average, a researcher would thus spend 8.1 hours analyzing a dataset with 300 T cells when only the tracks of crawling T cells (50%, i.e., 150 cells) are analyzed. If all T-cell tracks were to be analyzed, this time would further increase to 12.9 hours. Given that on average 10 datasets can be produced per day, manual analysis becomes a bottleneck, leading to delays in exhaustive data analysis and thus ultimately in research progress. The ^UFM^Track framework reduces analysis time by a factor of 3 when analyzing only crawling cells and 5 if all cell tracks are to be analyzed (Figure 5K).

With an average analysis time of 2.3 hours, the 10 experiments carried out in one day can be thoroughly analyzed within one day of machine time. This enables the scalability of flow-based immune cell migration experiments while simultaneously lifting the burden of tedious and time-consuming manual analysis from the researchers.

### V. Applicability of ^UFM^Track to different models of immune cell - endothelial cell interactions

To assess the broader applicability of the ^UFM^Track framework to different cell types and BBB models employed for the *in vitro* under-flow cell trafficking assays. To this end, we have performed the automated analysis of the multi-step T-cell migration cascade in a set of datasets (one to eight) across different cell types and assays. Specifically, we attempted to verify the feasibility of analysis of datasets where the cells forming the monolayer or the migrating cells have noticeably different appearances, cells from other species are used, specifically human-derived cells, and datasets with a different imaging modality. These include human peripheral blood mononuclear cells (hPBMCs), human T cells, bone marrow-derived macrophages (BMDM) interacting with a monolayer of mouse- and human-derived brain microvascular endothelial cells as well as with immobilized recombinant BBB adhesion molecules (See “Material and Methods” section for details). To assess the performance of the model in each condition, we have performed a visual inspection of the cell masks, transmigration masks, and cell tracks for few datasets in each assay type. We then analyzed cell behavior distribution or motility parameters relevant in each case.

#### 1. hPBMC on HBMEC monolayer

The interaction of hPBMCs with the BBB is altered under different disease conditions. We have therefore analyzed a dataset of human PBMCs on human brain microvascular endothelial cells (HBMEC) to quantify the cell behavior statistic. While the endothelial cells look significantly different than the pMBMECs, the segmentation and transmigration detection quality was sufficient to perform the cell migration analysis. This was also confirmed by the visual inspection of the cell and transmigration masks and cell tracking results (Figure 6 A). We then quantified the PBMC behavior distribution and the crawling and migration speed of the crawling PBMC (Figure 6 B, C).

**Figure 6.**
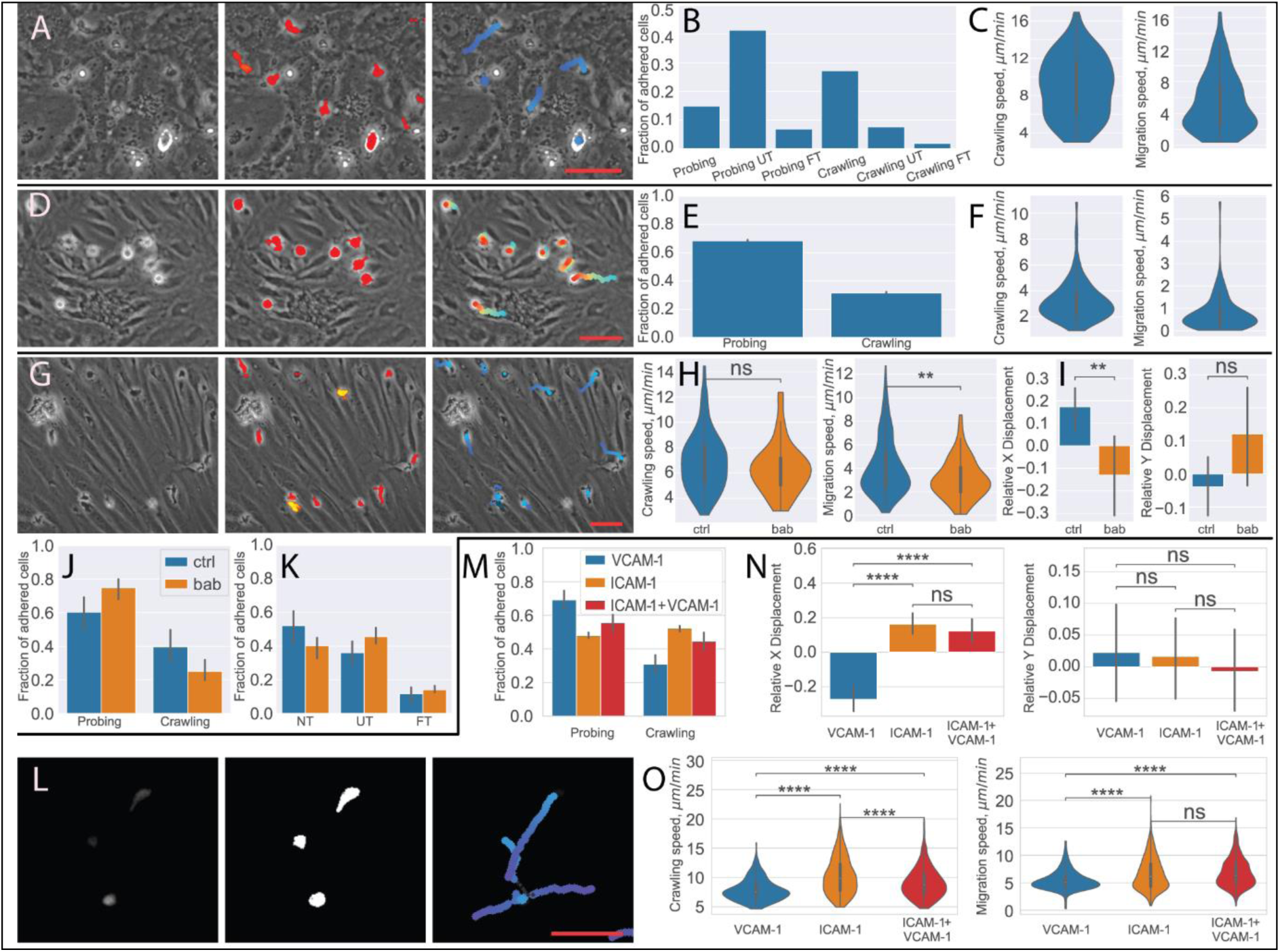
Analysis of different BBB models with ^UFM^Track. A-C. Human PBMCs interacting with HBMEC. A. Raw image frame, segmentation, and tracking results. B. PBMCs behavior distribution. C. Crawling and migration speed distribution. D-F. BMDM interacting with the pMBMEC monolayer. D. Raw image frame, segmentation, and tracking results. E. BMDM crawling vs probing behavior distribution. F. Crawling and migration speed distribution. G-K. Human T cells interacting with the EECM BMEC monolayer. G. Raw image frame, segmentation, and tracking results. H. Crawling and migration speed distributions for the control condition (ctrl) and with the blocking anti-bodies (bab). I. X and Y relative cell displacement. J. Crawling vs probing behavior comparison. K. Transmigrations behavior comparison. L-O. Human T cells interacting with immobilized recombinant BBB adhesion molecules. L. Raw fluorescent image frame, thresholding based segmentation, and tracking results. M. Crawling vs probing behavior distribution. N. X and Y relative displacement O. Crawling vs probing speed comparison. Scale bar 50 μm. Statistical tests are performed with the Mann-Whitney U test.

#### 2. BMDM on the pMBMEC monolayer

Invasion of macrophages into the CNS contributes to many neurological disorders. Therefore, we have explored if ^UFM^Track also allows us to analyze the interaction of BMDMs with the BBB under physiological flow in vitro. To this end, we have analyzed two datasets of mouse BMDMs interacting with pMBMEC monolayers (Figure 6 D). Despite the different behavior and appearance of the macrophages as compared to the T cells, the segmentation quality was sufficient to perform the cell tracking and analyze cell motility parameters (Figure 6 E, F), as confirmed by visual inspection of the macrophages masks and cell tracking results (Figure 6 D). Nonetheless, due to the different appearance of the flattening crawling cells and the transmigrated cells, the transmigration detection trained on the mouse T cells does not perform sufficiently well for the robust detection of transmigrating macrophages.

#### 3. Human T cells on EECM-BMEC monolayer

Next, we analyzed eight datasets of human T cells interacting with the human induced pluripotent stem cell (hiPSC) derived extended endothelial cell culture method brain microvascular endothelial cell (EECM-BMEC)-like cell monolayers (Figure 6 G). EECM-BMECs establish barrier properties comparable to primary human BMECs and display a mature immune phenotype allowing to perform studies for the first time in a fully autologous manner of immune cell interactions with patient-derived in vitro models of the BBB (35). Despite the cells being of different species, the T cells were successfully segmented, and their transmigration was detected, as confirmed by visual inspection of the T-cell and transmigration masks and cell tracking results (Figure 6 G).

We have confirmed the differences in migration statistics (Figure 6 J, K), migration and crawling speed of the crawling T cells (Figure 6 H), as well as the previously reported difference in relative (normalized to track pathlength) displacement (Figure 6 I) of the T cells treated with anti-α4-integrin + anti-β2-integrin blocking antibodies (bab) and control (ctrl) condition (36).

#### 4. Human T cells on immobilized recombinant BBB adhesion molecules

Finally, we have analyzed datasets acquired in a different imaging modality. Seven datasets of human T cells interacting with immobilized recombinant BBB adhesion molecules, namely ICAM-1, VCAM-1, and their combination (Figure 6 l). In this case, the fluorescently labeled cells were imaged with epi-fluorescent microscopy. In the absence of a monolayer, transmigration detection is not necessary. Thus we performed cell segmentation directly on the fluorescent signal (See Materials and Methods for details) using the same segmentation strategy as for the T-cell masks. The tracking was then performed the same way as for other experiments. The T cells were successfully segmented and tracked, as confirmed by visual inspection. We compared T cell behavior and motility parameters for three conditions of the recombinant adhesion molecules: ICAM-1, VCAM-1, and ICAM1 + VCAM1 (Figure 6 M, O). Consistent with previous observations (12,36) where the analysis was performed using the ImageJ (26) either manually or with the TrackMate (25) plugin, we observed that in the absence of ICAM-1, T cells were moving in the direction of flow (Figure 6 N). At the same time, ^UFM^Track allows for capturing the rapid accelerated motions of the cells and thus reconstructing longer continuous T cell tracks. This enables capturing the migrations properties of the consecutive steps of the T cell migration cascade on the call-by-cell basis.

## Discussion

In this study, we have developed the under-flow migration tracker (^UFM^Track) framework. It consists of independent modules and allows for segmentation (Figure 2), tracking (Figure 3), and motility analysis of immune cells, migrating on, across, and below (Figure 1D) the monolayer of brain microvascular endothelial cells under physiological flow in vitro. The developed method relies exclusively on phase-contrast imaging data (Figure 1B). Therefore, it does not require establishing fluorescent labels of the migrating immune cell population to be studied, avoids potential photo- toxicity to be considered for fluorescent imaging modalities (37–39), and is thus the preferred choice for analyzing the trafficking of sensitive cell types. T-cell segmentation and prediction of the transmigrated T-cell areas are performed using a custom 2D+T U-Net-like convolutional neural network. It enables reliable segmentation of T cells both above and below the pMBMEC monolayer. The existing particle and cell tracking toolkits consider migration of one cell type and thus are not suitable for the detection of distinct migration regimes and cell interactions (23–25,28).

Furthermore, to our knowledge, none of the existing algorithms consider migration under the flow causing rapid cell displacement. The tracking of T-cell interactions with the pMBMECs during all migration regimes under physiological flow required designing a new tracking algorithm considering rapid T-cell appearance, disappearance, and displacement in the field of view caused by the flow.

We have also developed approaches that resolve track intersections, i.e., identifying track segments corresponding to the same T cell before and after under-segmented track regions, which are inevitable during T-cell migration. By establishing the detection of T-cell crawling, probing, transmigration, and accelerated movement combined with reliable T-cell tracking, we have enabled the in-depth analysis of distinct migration regimes on a cell-by-cell basis. By reducing the dataset analysis time by a factor of 5, ^UFM^Track allows for performing a thorough analysis of 10 experiments that a researcher can carry out in one day within one day of machine-time. We have demonstrated that the automated analysis performs on par with manual analysis while improving accuracy and eliminating experimenter bias, and enables scalability of flow-based immune cell migration experiments by reducing the analysis cost and lifting the burden of time-consuming manual analysis.

In this work, we have demonstrated the applicability of the developed framework to the analysis of the multi-step extravasation of CD4^+^ and CD8^+^ T cells across non-stimulated or cytokine-stimulated pMBMEC monolayers under physiological flow. We have also demonstrated that the developed framework allows for automated analysis of T-cell behavior statistics and motility parameters of distinct T-cell migration regimes. Results of the automated analysis of CD4^+^ T-cell behavioral statistics performed with ^UFM^Track are in agreement with previous studies using manual data analysis (Figure 4B) (14). The automated analysis of datasets of CD8^+^ T-cell migration has shown comparable performance with manual analysis performed by one experimenter with four years of analysis experience and three less-experienced experimenters. At the same time, the variance of the results obtained manually showed significant experimenter bias that the automated analysis eliminated (Figure 5F). Additionally, the automated analysis allowed for in-depth analysis of all T-cell migration categories and precise evaluation of T-cell motility parameters, which was not achieved by manual T-cell tracking, even by the most experienced user.

We have further demonstrated the applicability of the ^UFM^Track framework to the analysis of different cell types (BMDM, PBMC), species (human-derived T cells, HBMEC, and EECM BMEC) as well as tracking and analysis of cell migration imaged using the fluorescent imaging modality. In all cases, the cells could be successfully segmented and tracked, and the cell migration analysis was performed on a cell-by-cell basis. The main challenge for the system is reliable segmentation of the T cells and the transmigrated T cells. In the case of HBMEC the monolayer has a dotted pattern unlike the rather smooth pMBMEC. This leads to more false positives in the T-cell segmentation at the level of individual frames, which nonetheless could be efficiently filtered at the level of cell tracking. In the case of the BMDMs, since they have significantly different appearances during transmigration from the T cells, reliable transmigration detection was not obtained. This prevents the system from quantifying the transmigration statistics of the BMDMs, yet the cell detection and thus motility parameter evaluation of BMDMs was successful (Figure 6 D-F). For all the other analyzed cell types, both the cell segmentation and transmigration detection model trained on the annotated CD4^+^ T cells also produced transmigration maps of sufficient quality for reliable transmigration detection.

The framework was able to successfully detect the cells of sizes that differ by almost a factor of two (Supplementary Figure 7). This demonstrates, that simply by discriminating cells by size, it is possible to extend ^UFM^Track to study interaction of several types of immune cells migrating on top of a cellular monolayer under flow and imaged by phase contrast.

Quantifying the fraction of transmigrated T cells is crucial for studying the molecular mechanisms governing the infiltration of autoaggressive T cells across the BBB into the CNS parenchyma and the immune surveillance. The role of specific endothelial or T-cell adhesion molecules in influencing the dynamic interactions of T cells with pMBMECs can, e.g., be probed by quantifying the ratio of T-cell crawling behavior to T-cell probing (Figure 6M) (14,40,41). The analysis procedure established in the ^UFM^Track framework is flexible and can be easily extended to evaluate further properties of the migrating cells. For example, the number of times a T cell interrupts its crawling regime by switching to short probing behavior on the endothelium can be evaluated, thus providing information on the distribution of “hot spots” on the endothelium. Additionally, experiment scalability and the analysis on a cell-by-cell basis enable the search for distinct populations of T-cells, e.g., according to the distribution of probing to crawling behavioral ratios. The strength of T-cell adhesion to the endothelium can be probed with the developed framework using the measure of T-cell detachment rate and the distribution of the previously overlooked T-cell accelerated movement occurrences and accelerated movement speed. Finally, the possibility of quantifying the motility parameters of the transmigrated T cells enables future studies involving multilayer in vitro BBB models that include the vascular basement membrane in addition to mural cells such as pericytes and astrocytes mimicking the entire neurovascular unit.

While in this work we have focused primarily on the analysis of mouse T cells interacting with pMBMEC under flow or other immune cells interacting with the endothelial monolayer, the methods we present establish a foundation for a broad range of studies involving in vitro under-flow studies of immune cell trafficking. The tracking algorithm and motility analysis developed in the ^UFM^Track framework are also directly applicable to the analysis of immune cell interactions with recombinant BBB adhesion molecules under flow. In this case, either phase-contrast or epi-fluorescent imaging can be employed when studying fluorescently labeled immune cells, as demonstrated. The T-cell segmentation based on deep neural networks can be applied to studies of the trafficking of other immune cell subsets on and across the pMBMEC monolayer. We demonstrated the successful application of the ^UFM^Track to cells with significantly different appearances, such as the HBMEC cellular monolayers.

We make UFMTrack available to the community to build upon it for addressing their specific research questions. The modularity of the framework simplifies the adaptation to analysis of different cell types. The application to immune cell trafficking across other endothelial monolayers, including those from different species or vascular beds as well as lymphatic endothelial cells where the cell appearance is drastically different from the ones demonstrated here, is possible but requires fine-tuning of the trained segmentation model. While training the models presented here required a large dataset of annotated T-cell masks, and transmigration masks, this can be largely avoided for future development. Future models can be developed more efficiently by leveraging our existing annotated dataset of T-cell migration on the pMBMECs while employing transfer learning approaches in a multitask framework with weak supervision and fluorescent labels as auxiliary learning targets to adapt the model for the segmentation of cells with different appearances. This can be further improved by adopting self-supervised contrastive or masked learning methods, which have demonstrated significant advancements in model pretraining in recent years (42–45). This is the subject of our future studies. The framework could be employed in combination of other methods, e.g. by incorporating fluorescently labeled endothelial adherens junctions one could differentiate the trans- from the paracellular transmigration in the migrating cells. In this case the transmigration locations detected by the ^UFM^Track framework could be automatically extracted for analysis. One could employ the tracking under flow implemented in the ^UFM^Track framework for tracking the immune cells migrating on the vascular wall *in vivo* to capture the accelerated movement of cells. By sharing the data and open-source code of the ^UFM^Track framework, i.e., the training data used for the segmentation model training, trained models, as well as the model architecture, and full under-flow T-cell tracking and migration analysis pipeline, we hope to encourage the community to pursue these developments to advance the field.

One current limitation of the method is the performance reduction of the T-cell tracking when the density of migrating T cells significantly increases (>250 cells/dataset). Thus, the recommended T cell concentration is about 100-150k/ml. However, to avoid non-physiological interactions between migrating T cells and T cell clumping, the T-cell density should be kept at moderate levels anyway.

Additionally, analyzing data from 8 stitched FoVs allows for obtaining comparable statistics. The current analysis of the behavioral statistics quantifies the fraction of different T-cell migration regimes with respect to the number of adhered cells instead of the total T-cell count passing through the microfluidic device. While the number of fast-moving T cells during the accumulation phase cannot be directly counted using the imaging modality employed here, it can be estimated indirectly from the imaging data with additional calibration experiments and a dedicated machine learning model. Another limitation is that currently, we do not consider dividing immune cells. While cell division events happen rarely (<1% of T cells) and are not primary events for the study of immune cell interaction with the BBB model, the modular architecture of our framework will facilitate future extension to detect cell division.

By enabling experiment scalability, unbiased analysis with advanced accuracy, and an in-depth analysis of large datasets of T-cell dynamics under flow, the computational and analytical framework presented here contributes to the 3R principle when studying the interaction of cells derived from animal models by reducing the number of animals to be sacrificed. ^UFM^Track can be employed for fundamental research of the molecular mechanisms governing immune cell trafficking across a range of vascular beds, screening of pharmaceutical treatments, as well as for personalized medicine based on the evaluation of treatment efficacy on patient-derived T-cell migration behavior on patient-derived endothelial monolayers. One can furthermore envisage that ^UFM^Track can be suitable to study cancer cell metastasis across different endothelial cell monolayers. Eventually, the developed framework can be extended to real-time operation during image acquisition. Combined with transgenic photo-convertible immune cells allowing for the photoconversion of immune cells according to their behavior will allow for subsequent fluorescent cell sorting and scRNA-Seq analysis. Such advanced studies are needed to reveal the role of genetic differences governing T-cell migration regimes.

## Materials and Methods

### 1. pMBMEC cell culture

Primary mouse brain microvascular endothelial cells (pMBMECs) were isolated from 8-12 weeks old C57BL/6J WT mice and cultured exactly as described before (12,46). Intact monolayers were stimulated or not with 10 ng/mL of recombinant mouse TNF, 20 ng/mL of recombinant mouse IL1-β for, or 5ng/mL recombinant mouse TNF + 100 U/mL recombinant mouse IFN-γ 16-24 hours prior to the assays as previously described (32).

### 2. Human brain microvascular endothelial cells

Human brain microvascular endothelial cells (HBMEC) were kindly provided by Prof. Nicholas Schwab (University of Münster, Germany). They were cultured to confluency on speed coating solution coated µ-Dishes (35 mm, low, iBidi) for 2 days as previously described (32).

### 3. Extended endothelial culture method—brain microvascular endothelial cell-like cells

The Extended Endothelial Cell Culture Method (EECM) was used to differentiate human induced pluripotent stem cells (hiPSCs) to brain microvascular endothelial cell (BMEC)-like cells were employed as a human in vitro model of the BBB. In brief, hiPSCs from one healthy control (HC) (cell line ID: LNISi002-B) were established from erythroblasts in the laboratory of Renaud DuPasquier (University of Lausanne, Switzerland). EECM-BMEC-like cells were cultured to confluency on collagen IV and fibronectin-coated µ-Dishes (35 mm, low, iBidi) for 2 days (36).

### 4. T cell and hPBMC preparation

Naïve CD4^+^ and CD8^+^ T-cell isolation: Peripheral lymph nodes and spleens from 2D2 and OT-I C57BL/6J mice were harvested, and single-cell suspensions were obtained by homogenization and filtration through a sterile 100 μm nylon mesh. A second filtration was applied after erythrocyte lysis (0.83% NH4Cl, Tris-HCl). 2D2 and OT-I cells were isolated respectively with magnetic CD4^+^ and CD8^+^ T cell selection beads (EasySep, STEMCELL Technologies).

In vitro activation of naïve CD8^+^ T cells: OT-I CD8^+^ T cells were activated as described before (13,41). Activated CD8^+^ T cells were cultured in IL-2-containing media for 3 days post-activation.

In vitro activation of naïve CD4^+^ T cells: 2D2 CD4^+^ T cells were activated as described before (32).

Activated CD4^+^ T cells were cultured in IL-2-containing media for 24 additional hours.

hPBMC were isolated from buffy coats of healthy donors by Ficoll-Paque™ Plus (Cytiva) density gradient and were frozen and stored in a liquid nitrogen tank until use as previously described in (36).

Human CD4^+^ T cells were isolated by employing a CD4^+^ T cells isolation kit (Miltenyi Biotec kit) following the provider’s instructions. Subsequently, effector/memory CD4^+^ T cells were sorted by Fluorescence-Activated Cell Sorting (FACS) into different Th subsets according to their specific surface expression of chemokine receptors (CCR6^−^CXCR3^+^CCR4^−^ for Th1; CCR6^+^CXCR3^+^CCR4^−^ for Th1*; CCR6^−^CXCR3^−^CCR4^+^ for Th2; CCR6^+^CXCR3^−^CCR4^+^ for Th17) and expanded as previously described (1,33,40,47–49).

### 5. BMDM preparation

BMDMs were prepared as described in (Berve et al., manuscript in preparation). In brief, the mouse bone marrow was harvested and washed through a 100 µm nylon mesh. After 7 days of cell culture were plated in a density of 17-20 × 10^6^ cells/mL in BMDM medium supplemented with 5ng/mL of recombinant mouse macrophage colony stimulating factor (mCSF, R&D Biosystems,

Minneapolis, USA, 416-ML-500) onto non-treated 100mm tissue culture Petri dishes (Greiner Bio-One, St. Gallen, Switzerland) for 7 days at 37°C and 5% CO2 the cells were harvested by incubation with 0.05% Trypsin.

### 6. T cell binding assays under physiological flow conditions

For live-cell imaging of T-cell interaction with recombinant BBB cell adhesion molecules under the physiological flow condition, µ-Dishes (35 mm, low, iBidi) were coated with 1.54 μg/mL human recombinant VCAM-1 (Biolegend) and/or 1.14 μg/mL human recombinant ICAM-1 (R&D system) in DPBS for 1 h at 37°C and blocked with 1.5% (v/v) bovine serum albumin (Sigma-Aldrich) in DPBS overnight at 4°C.

CMFDA prelabelled Th cells were incubated with the isotype control antibody (30 μg/mL) for 30 minutes at 37°C (8% CO2) prior to imaging (exact antibody concentrations are illustrated in the figure legend). The cells were subsequently perfused on top of the pre-coated dishes at a concentration of 10^6^/mL in MAM as previously described (12,36).

### 7. In vitro under-flow T-cell migration assay

We studied the multi-step T-cell migration across monolayers of primary mouse brain microvascular endothelial cells (pMBMECs) in a custom made microfluidic device under physiological flow by in vitro live-cell imaging according to the previously established procedure (14,32). The custom-made flow chamber was described in depth before (11,46). The flow chamber was placed on top of the pMBMECs, previously cultured in Ibidi µ-Dish to confluency, and connected to the flow system filled with migration assay medium (MAM) (12). During the accumulation phase, T cells at a concentration between 55k/mL and 166k/mL were perfused on top for 5 minutes under low shear stress of 0.1 dynes/cm^2^, allowing them to settle on top of the pMBMEC monolayer. We used lower T-cell concentration compared to previous studies to enable automated T-cell interaction analysis. Afterward, the flow was increased to physiological levels with a flow shear stress of 1.5 dynes/cm^2^ to study post-arrest T-cell behavior on pMBMECs for 27 minutes (Figure 1A, B).

### 8. Data acquisition

The timelapse imaging was performed during both accumulation and physiological flow phases using phase-contrast imaging at a framerate of 6 frames/minute or 12 frames/minute, subsampled to 6 frames/minute and resolution of 0.629 µm/pixel with a 10x objective. In this modality the acquired images are gray scale, and the T cells, especially after the migration across the pMBMEC monolayer, have a similar appearance to the pMBMECs, making the T-cell segmentation task very challenging (Figure 1C). The data was acquired in tiles of 870 × 650 µm^2^ (1389 × 1041 pixels) with an overlap of 100 µm, leading to a total acquired image area of 3170 × 1220 µm^2^. For each experiment, the dataset consists of 30 timeframes of T-cell accumulation and 162 timeframes of dynamic T-cell interactions with the pMBMEC monolayer under physiological flow. The timestep between sequential timeframes for each tile is 10 s. The total acquired area is thus limited by the acquisition speed or by the flow chamber size.

The HBMEC dataset was acquired as previously described (36). In brief, 2 tiles were acquired of total size 3891 × 1504 pixels with pixel size 0.69 µm and framerate 6 frames/minute. The accumulation phase was 4 minutes, and the physiological flow phase was 5 minutes.

The BMDM dataset was acquired similarly to the T cell interacting with the pMBMEC datasets. In brief, one tile was acquired of size of 2048 × 1504 or 1388 × 1040 pixels with pixel size 0.69 or 0.629 µm and framerate 6 frames/minute. The accumulation phase lasted 6 minutes, and the physiological flow phase was 25 minutes.

The EECM-BMEC dataset was acquired as previously described (36). In brief, 2 tiles were acquired of total size 3892 × 1504 or 2638 × 1048 pixels with pixel size 0.69 or 0.629 µm and framerate 6 frames/minute.

The dataset of human T cells on immobilized recombinant BBB adhesion molecules was acquired as previously described (36). In brief, 2 tiles were acquired of total size 3891 × 1504 pixels with pixel size 0.69 µm and framerate 6 frames/minute. The accumulation phase was 4 minutes, and the physiological flow phase was 5 minutes. In this dataset, there is only one compartment, thus excluding any transmigration of the T cells. We, therefore, have employed the preprocessed fluorescent images instead of the cell probability map and transmigration map produced by the segmentation neural network. In this case, the centroids are defined as local maxima localized on the preprocessed fluorescent images. The fluorescent images were preprocessed first by performing Gaussian blur filtering in 2D with a standard deviation of 0.75. Next, thresholding was applied to eliminate the background noise, clipping high brightness values (color values with abundance <0.3% of threshold abundance), scaling to 0-255 8 bit, and saving in a lossless format. Finally, the 2 tiles were stitched together by matching the integrated color distribution in the X direction in the overlapping area.

### 9. Manual analysis of T-cell migration

Manual cell analysis and tracking were performed according to the previously established procedure using ImageJ software (ImageJ software, National Institute of Health, Bethesda, MD, USA) (13). The number of arrested T cells was thus counted at timeframe 33 of the subset. The behavior of arrested T cells was defined and expressed as fractions of arrested T cells set to 100% as follows:

- T cells that detached during the observation time (“detached”)
- T cells that migrated out of the FoV detached during the observation time (“out of FoV”)
- T cells that continuously crawled on the pMBMEC monolayer (“crawling”)
- T cells that remained in the same location (displacement less than twice the cell size) while actively interacting with the pMBMEC monolayer (“probing”)
- T cells that crossed the pMBMEC monolayer with or without prior crawling (“crawling full transmigration” and “probing full transmigration”). The event of T-cell transmigration across the pMBMECs monolayer became obvious due to the change of appearance of the transmigrated part of the T cells from phase bright (on top of the pMBMECs monolayer) to phase dark.
- T cells that partially crossed the pMBMEC monolayer, then retracted the protrusions and continued to migrate above the monolayer (“crawling uncompleted transmigration” and “probing uncompleted transmigration”).

### 10. ^UFM^Track performance evaluation

The inference of T cell masks and transmigration masks was performed on a dual-CPU Intel(R) Xeon(R) CPU E5-2670 v3 @ 2.30GHz, 256GB RAM node equipped with 8 Graphical Processing Units (GPU) NVIDIA GeForce GTX TITAN X / GeForce GTX 1080. Frame alignment and segmentation were performed on an Intel(R) Core(TM) CPU i7-4771 @ 3.50GHz, 32GB RAM workstation with NVIDIA GeForce GTX TITAN GPU. Cell tracking was performed on a dual-CPU Intel(R) Xeon(R) CPU E5-2643 v2 @ 3.50GHz, 256GB RAM workstation.

### 11. Performance analysis

The statistical tests are performed with the Mann-Whitney U test. Performance of the segmentation models was evaluated based on the pixel classification metrics evaluated using the scikit-learn library (50). Specifically, the Average Precision is evaluated on the test set as the area under the Precision-Recall curve, as the weighted mean of precisions achieved at each threshold, with the change in recall from the previous threshold used as the weight. The segmentation thresholds for the cell mask and the transmigrated cell mask predictions were selected to maximize the F1 score.

## Acknowledgments

We thank Kristina Berve for providing BMDM datasets for analysis with the ^UFM^Track. We owe sincere thanks to Dr. James McGrath and Danial Ahmad (Rochester University, NY, USA) for their detailed and insightful feedback on our manuscript. This work was supported by an UniBe ID Grant to MV, BE, and AA and the Microscopy Imaging Center (MIC) of the University of Bern.

## Data Availability

The source code for the T-cell segmentation models, data preprocessing, training, performance evaluation, and inference scripts are available on GitHub at https://github.com/neworldemancer/UFMSegm. This repository also contains win64 binaries used for the watershed-based segmentation of the predicted T cell probability maps and transmigration probability maps.

The source code for the under-flow T-cell tracking and migration analysis, along with the scripts used for performance evaluation of the framework, is available on GitHub at https://github.com/neworldemancer/UFMTrack.

The training phase-contrast data with manual annotations, the reference datasets for histogram normalization, and trained models are available on Zenodo: [http://doi.org/10.5281/zenodo.7489557].

The datasets used for evaluation of our framework are available on Zenodo: [http://doi.org/10.5281/zenodo.7489972, http://doi.org/10.5281/zenodo.7489984, http://doi.org/10.5281/zenodo.7489996, http://doi.org/10.5281/zenodo.7490012, http://doi.org/10.5281/zenodo.7490020, http://doi.org/10.5281/zenodo.7490029].

## Authors contribution

M.V. implemented the ^UFM^Track software framework, conducted automated analysis and performance evaluation, and contributed to experimental design and imaging.

M.V. and B.E. wrote the manuscript with contributions from S.S., S.A., and L.M.

M.V., with contributions from S.A., L.M., and S.S., conceptualized the T-cell migration analysis pipeline.

L.M., S.A., and S.S. conducted wet lab experiments and imaging.

S.A., L.M., A.M., and M.V. performed annotation of the datasets for the segmentation model training.

S.A., L.G., A.P., and M.V. performed manual data analysis.

A.A. and B.E. designed the research together with M.V., acquired funding, and coordinated the collaboration.

