## Supplementary information for "^UFM^Track: Under-Flow Migration Tracker enabling analysis of the entire multi-step immune cell extravasation cascade across the blood-brain barrier in microfluidic devices"

### 1. ***Segmentation models***

#### *Training data*

Deep neural networks achieve high performance but require a large amount of data for model training. Thus, in addition to preparing a large high-quality training dataset of annotated T cells, we employed additional learning targets and data augmentation to improve model performance. To train the models, we have manually annotated subsets from 4 independent experiments, combined corresponding to approximately 30 min of time-lapse acquisition of 880 × 660 $\mu m^2$, 226 Megapixels in total. The data was split into non-adjacent training and validation datasets of 154 and 37 Megapixels correspondingly. We have annotated the masks of whole T cells ("T cell mask") and the mask of the transmigrated part of the T cells ("transmigration mask"). Annotation masks were normalized to have the values 0 and 1. Additionally, the center points of the touching and overlapping T cells were annotated, while for the rest of the T cells, the center points were obtained as the center of mass of the T cell mask. Afterwards, maps of centroids were generated where each centroid was represented by higher values in the map following a 2-dimensional Gaussian function with width $\sigma = \frac{1}{10} r_{cent}$ where $r_{cent}$ is the distance to the nearest centroid. This width was then clipped to the range $\sigma \in [1.5, 4]$ pixels so that all centroids have a comparable scale. To prioritize the T cell segmentation quality in the vicinity of the transmigrated part of the T cells, we created distance maps to the nearest transmigrated T cell pixel $r_{tm}$. We then created a weight map as $w_{cell} = 1 + \alpha_w \exp\left(-\frac{r_{tm}}{d_{tm}}\right)$. The two annotation masks, the centroids map, and the weight map (Figure 2) were used in the model loss function (see Training section below).

Consecutive frames of the time-lapse phase-contrast imaging datasets were aligned by a cross-correlation and global offset optimization analogous to the one described in Vladymyrov et al. (1). Additionally, we have applied histogram normalization of the phase-contrast images (2) to achieve a coherent brightness distribution across all datasets. During histogram normalization, we adjusted the brightness for all pixels such that the brightness distribution of the pMBMECs monolayer in each image matched the brightness distribution of the pMBMECs monolayer in a reference image. We have selected one dataset as a reference according to the best validation performance of trained models. Then, following common practice, we have standardized the brightness distribution according to brightness values between the $2.5^{th}$ and $97.5^{th}$ percentile.

During training, we performed data augmentation consisting of random rotation by arbitrary angle, brightness change (<50%), contrast change (<50%), and affine deformations. During rotation and deformation transformations, the image and all label masks were transformed coherently, employing bilinear interpolation for the input image, centroids, and the weight map. For the T cell masks and the transmigration masks, the "nearest" interpolation was applied. At each training iteration, a new random transformation was applied to a randomly selected crop of the training dataset, thus producing an unlimited number of possible variations.

*Training*

We trained the model in a multitask learning framework by optimizing the loss function

$$L = w_{cell}\, L_{cell} + L_{tm} + \alpha\, L_{cent},$$

where $w_{cell}$ is the weight map prioritizing T cell segmentation quality in the vicinity of the transmigrated part of the T cells. Here

$$L_{cell} = -\Sigma_{\mathrm{b}}(\mathrm{l_c} \log \widehat{p_c} + (1 - \mathrm{l_c})\log(1 - \widehat{p_c}))/\mathrm{n_b}$$

is the mean cross-entropy for the T cell mask predicted probability $\widehat{p_c}$ given the ground truth label $l_c$;

$$L_{tm} = -\Sigma_{\mathrm{b}}(\mathrm{l_{tm}} \log \widehat{p_{tm}} + (1 - \mathrm{l_{tm}})\log(1 - \widehat{p_{tm}}))/\mathrm{n_b}$$

is the mean cross-entropy for the transmigrated T cell mask prediction probability $\widehat{p_{tm}}$ given ground truth transmigration label $l_{tm}$;

$$L_{cent} = \Sigma_{\mathrm{b}}(|\widehat{\mathrm{h_{cent}}} - l_{cent}|)/n_b$$

is the $L_1$distance between the predicted value on the centroids map $\widehat{\mathrm{h_{cent}}}$ and corresponding label $l_{cent}$. The summation was performed over all $\mathrm{n_b}$ pixels in the training batch.

Coefficients $\alpha = 5$, $\alpha_w = 2$, and the characteristic distance from the nearest transmigrated T cell $d_{tm} = 15\ \mu m$ were chosen empirically to maximize the Intersection-over-Union (IoU) metric value in the validation.

The models were trained on image crops with the size of 265 × 256 pixels, using Adam optimizer (3) with a batch size of 2, for 40k iterations (~840 epochs). The training took 17 hours for 2D and 29 hours for the 2D+T models using a single GeForce GTX TITAN X GPU. The learning rate was set to $1.5 \times 10^{-3}$. During training, the learning rate was adjusted in stages. It was ramped up starting $1/4$ of the nominal value for 1k iterations and then halved after 20k iterations, as shown in Supplementary Figure 1A. The training was stopped when the validation IoU metric reached its maximum for the T-cell mask prediction. The loss evolution along the training is shown in Supplementary Figure 1B.

*Performance evaluation*

Next, we evaluated the pixel-wise performance of the trained T-cell segmentation models for prediction of the T-cell mask and the transmigration mask on the validation dataset. We assessed the F1 Score, Jaccard index, and average precision (AP), as shown in Supplementary Figure 2 and Table 1 for the T-cell and transmigration masks for both the 2D vs 2D+T models. For the T-cell mask prediction, the 2D+T model outperformed the 2D model by a notable 9%. At the same time, for the transmigration mask, which is much more difficult to detect, the 2D model performance reached only 54%, which is insufficient for the reliable detection of T cells migrating across the pMBMEC monolayer. In this task, the 2D+T model outperformed the 2D model by 32% AP. We observed that the 2D+T model was sensitive to frame misalignment, leading to false positive detection of transmigrated T cells. Thus, to generate the preliminary T-cell masks used for frame alignment and histogram normalization, we employed the 2D model. Afterwards, for the T cell segmentation and transmigration detection, we employed the 2D+T model.

*Image data processing*

For T-cell segmentation we followed a procedure similar to the one used for training data preprocessing. We first used the 2D model to localize T cells in individual image frames. We then dilated the predicted T cell masks by 7 pixels, such that the bright halo on the phase-contrast images surrounding the T cells arresting and migrating on top of the pMBMEC monolayer was enclosed in the masked region.

Next, we performed image histogram normalization of the image frames based on the brightness distribution of the pMBMECs. For this we used the dilated T cell masks obtained in the previous step. The same histogram reference dataset previously used for the model training was applied. Histogram normalization needed to be performed separately for the image frames obtained during the accumulation phase and the physiological flow phase of the assay. This is due to the overall image brightness increase after increasing the flow rate due to slight change of the sample position.

The histogram-normalized image sequence was then aligned based on the pMBMECs, using the masked imaged with the same dilated T cell mask. Frame alignment (1) was performed by 5 crops the size of 512 × 512 pix each in the corners and in the middle of the frame. We first measured offsets $dr_{c,i\to j}$ and offset errors $\sigma_{c,i\to j}$ between pairs of image frame crops $c$ at timepoints $i$ and $j$, $\forall i,\ j = i..i+10$. Then we employed global solving to find the absolute position $r_{c,i}$ and the respective position errors $\sigma_{c,i}$ for each crop $c$. Finally, we obtained the absolute positions of the entire frames by weighting the positions obtained by the crops: $r_i = \frac{\sum_c \frac{r_{c,i}}{\sigma_{c,i}}}{\sum_c \frac{1}{\sigma_{c,i}}}$.

The sequence of histogram-normalized and aligned image frames was then fed into the trained 2D+T segmentation model to obtain the T cell and transmigration probability maps, as well as the T cell centroids. To this end we processed the image sequences in patches of 512 × 512 × 5 pixels, with an overlap of 183 pixels. The size of the receptive field of our T cell segmentation model governed this overlap. Notably, while the models were trained on image crops of 256 × 256 pixels, the fully convolutional architecture allows the inference to be performed on any input size. Input size is thus limited solely by the available GPU memory.

The separate tiles of the tiled image sequence acquisition were aligned by overlapping parts of the raw image data. Predicted maps were then stitched together, where maximum predicted probability values were taken for overlapping tile regions.

To perform T-cell segmentation, we obtained T-cell masks from the T-cell probability map by applying a threshold of 0.33. Seed points were obtained from the T cell centroid map with a threshold of 0.16. These threshold values maximized the F1 score of the T cell mask prediction, as seen in Supplementary Figure 2. The segmentation was then performed with a watershed algorithm based on the T-cell mask and the seed points. A stitched phase-contrast image sequence can be seen in Supplementary Video 1 and overlayed with the segmented cells and highlighted transmigration mask in Supplementary Video 2.

### 2. T-cell tracking

*Linking T cells*

As the first step, we identified all possible connections of T cells between the timeframes. We constructed a graph representation of the datasets, such that graph nodes corresponded to the T cells at all timeframes, and the edges in the graph corresponded to links connecting the nodes adjacent in space and time. Along the tracking procedure, we then sought to identify links connecting the same T cell over time. In cases where T cells touched each other, i.e., when several centroids were detected within the same T cell mask or adjacent T cell masks with distances below 10 µm (i.e., potentially over-segmented cells), we considered them as one node. At this step, we accounted for this under-segmentation by introducing the node multiplicity number $m$ equal to the number of constituting components. These adjacent cells were assigned to separate tracks after the reliable track segments of isolated T cells were constructed (see Resolving global track consistency section below).

The links between T cells were searched within a radius $R = dt \cdot 15\ \mu m + 10\ \mu m$, where time difference dt is between 1 and 3 timeframes (Figure 3C).

*Track segment search*

This step aimed to find continuous track segments of T cells or under-segmented groups of T cells crawling on top of or below the pMBMEC monolayer without accelerated movement segments on the T cell track characterized by rapid T cell displacements. The whole dataset of T cells across all timepoints was represented as a graph. Each vertex corresponds either to a T cell (multiplicity m=1) or a group of potentially under-segmented T cells (m>1). The vertices are connected according to links obtained at the linking step. Track segments were found by performing global optimization to find consistent connectivity of vertices across the timepoints by employing an approach inspired by the conservation tracking algorithm summarized in Schiegg et al. (4). Each vertex was represented as a set of incoming (Left, L) and outgoing (Right, R) links, and additional “not connected” (NC) left and right links (Figure 3C, D, Supplementary Figure 3A, Supplementary Table 3). The optimization procedure selected the links that were most likely to be connections between the same T cell in different timeframes. The total number of selected links for each vertex on the left and right sides was limited to vertex multiplicity m. Optimization was performed using the CP-SAT constrained optimization procedure using the open-source OR-Tools library (5).

We employed the following constraints for each node.

Constrain “not connected” variable:

$$c_{nc\_l} = \begin{cases} 0 & \text{if } \sum_i c_l[i] > 0 \\ 1 & \text{otherwise} \end{cases}$$

Limit total “multiplicity” of node connections:

$$1 \leq c_{nc\_l} + \sum_i c_l[i] \leq m$$

Link the “selected” flag of a connection with its multiplicity:

$$c_{w\_l}[i] = \begin{cases} 1 & \text{if } c_l[i] > 0 \\ 0 & \text{otherwise} \end{cases}$$

And similarly for the right side.

If the node has connection candidates on both sides, then evaluate the multiplicity difference:

$$d_{abs\_lr} = \begin{cases} 0 & \text{if} \quad \sum_i c_r[i] == 0 \quad \text{or} \quad \sum_i c_l[i] == 0 \\ \left| \sum_i c_r[i] - \sum_i c_l[i] \right| & \text{otherwise} \end{cases}$$

On each edge between connected nodes, the multiplicity on both sides should be the same:

$$c_{r,left\ node}\ [i_{connection\ to\ right\ node}] == c_{l,right\ node}[j_{connection\ to\ left\ node}]$$

Then the loss function for each node is

$$L_n = d_{abs\_lr} * w_m + \sum_{l,r} c_{nc}\ * w_{nc}\ + \sum_i c_{w\_r}[i] * w[i]$$

This loss has three terms. The first one penalizes differences between connection multiplicity on the left and right sides. The second penalizes solutions where the node is not connected on either side. The last term weights each selected connection to a node on the right. Here $w[i]$ is the weight of the connection to the corresponding right node. Importantly, connection weighting is independent of the connection multiplicity. This design choice was made explicitly since node multiplicity estimates available at this stage are preliminary.

To account for T cell detection inefficiency at the edge of the FoV, the penalty for not connected nodes $w_{nc}$ is attenuated close to the FoV edge. To this end the weight of a not-connected node exponentially decays when the T cell is closer than $d_0 = 20\ \mu m$ to the edge:

$$w_{nc}\ = f_{edge}\ * w_{nc_0}\ ,$$

where

$$f_{edge}\ = \begin{cases} 1 & \text{if} \quad d > d_0 \\ \exp\left(\frac{d_0 - d}{d_c}\right) & \text{otherwise} \end{cases}$$

$d_c$ is chosen such that $w_{nc} = w_{nc\_0}$ if $d = d_0$ and $w_{nc} = 1$ if $d = r_{cell}$, where the T cell radius (half of the crawling T cell length) $r_{cell} = 10\ \mu m$.

The constants $w_m = 3$, $w\_nc = 9$ were chosen empirically to achieve reliable reconstruction of long tracks.

Connections between nodes $c_1$ and $c_2$ at timeframes $t_1$ and $t_2$ correspondingly was considered if they are at most three timeframes apart $1 \leq t_2 - t_1 \leq dt_{max}\ = 3$, according to the output of the linking step. The connection weight $w$ between these nodes was evaluated as the negative log-likelihood (NLL) of the connection between these nodes:

$$w = w_v + w_A + w_{dt}$$

$w_v = \frac{1}{2}\left(\frac{v - v_\mu}{\sigma_v}\right)^2$ – is the NLL of the crawling speed (path length over time), where $v = \frac{r_{12}}{(t_2 - t_1) * \Delta t}$ is the speed estimate based on the positions $r_1, r_2$ of the closest of the T cells constituting nodes $c_1$, $c_2$. $\Delta t = 10s$ – is the time step between consecutive timeframes. $v_\mu = 9\ \mu m/min$ and $\sigma_v = 8.5\ \mu m/min$ are prior population mean and standard deviation of T-cell crawling speed.

$w_a = \frac{1}{2}\left(\frac{\Delta A}{\sigma_A}\right)^2$, where $\Delta A = \min_{cells\ in\ c_1, c_2} 2\frac{A_1 - A_2}{A_1 + A_2}$, $A_1$ and $A_2$ are the areas of the detected constituting cells of nodes $c_1$, $c_2$. $\sigma_A = 0.397$ is the prior standard deviation of $\Delta A$ estimated from the data.

$w_{dt} = w_{missing} * (t_2 - t_1 - 1)$ – penalizes connections between non-consecutive timeframes while accounting for possible T-cell detection inefficiency. The constant $w_{missing} = 2.3$ was chosen empirically.

To reduce the time required for this optimization, the procedure was performed independently for each connected by links components of the dataset graph.

*Resolving global track consistency*

In this step, the scope of T cell tracking is shifted from individual nodes (representing T cells at particular timeframes and groups of under-segmented T cells) to the track segments – unambiguous sequences of nodes and vertices at the endpoints of the segments. These are track start and end points, points of merging and separation of track segments in case of under-segmentation, as well as ambiguous points on a track. The latter were identified by sudden T cell displacement, a hallmark of detaching and reattaching T cells and T cells transitioning from properly segmented to under-segmented T cells or vice versa (Figure 3D, E).

First, the segments were searched as a sequence of links between nodes connected to only one node on the following timeframe or nodes branching to multiple nodes and later rejoining to one node. This corresponded to over-segmentation at some points of the track or an under-segmented group of T cells shortly separating to being properly segmented before re-joining. These branching and re-joining node sequences were considered one segment at this stage. The vertices were the starting and end nodes of the segments, including the branching points between the segments. This significantly simplified the graph representation of the T-cell tracks. Next, the segments were split according to additional criteria, such as large displacements caused by T cell detachment or intersection with another T cell. Empirically we have chosen the displacement threshold for splitting a segment to be $d > d_{thres} = \mu_d + 0.5\ \sigma_d$, where $\mu_d$, $\sigma_d$ – are the mean and standard deviation of T cell displacements between consecutive timeframes along the given segment. The node with the larger area was assigned as the new vertex. With each segment we associated the segment multiplicity $M$. Intuitively, this corresponds to the conservation of T cell number along a track segment. At first, it was estimated as $M_0 = \underset{t}{Median}(\sum_{cells} m)$ – median multiplicity of constituting nodes along the track (Figure 3D).

Next, we searched for potential missing track segments due to accelerated T cell movement under flow and missing links in the track crossing points. Both are characterized by considerable displacement length such that they were not detected during the linking step. Thus we will refer to both as "jumps" in this section. The two categories differ in the distribution of displacement length and direction. To perform this search, the likelihood $w_{ps}$ of the potential segments between vertices according to the average T cell "jump" length and direction for displacement caused by flow and T cell mask fusion due to under-segmentation was evaluated. In the case of jumps caused by under-segmentation, the square root of the displacement is approximately normally distributed. Thus, the negative log-likelihood of a potential segment was estimated as

$$w_{sj0} = c_{sj} \, \frac{1}{2}\left(\frac{\sqrt{d} - \mu_{sj}}{\sigma_{sj}}\right)^2 ,$$

where $\mu_{sj}, \sigma_{sj}$ – are the mean and standard deviation of the distributions of the square root of the displacement $\sqrt{d}$ and

$$c_{sj} = \begin{cases} 1 & \text{if } \sqrt{d} < \mu_{sj} \\ 2 & \text{otherwise} \end{cases}$$

(see Supplementary Figure 4A). The time duration of this potential segmentation-caused jump segment is also penalized, leading to the total NLL:

$$w_{sj} = \frac{1}{\sqrt{2}}(w_{sj0} + c_{sj,t}(dt - 1)),$$

where $dt$ is segment duration in timeframes and $c_{sj,t} = 1$.

For the T cell jumps caused by flow, we empirically estimated (see Supplementary Figure 4B-D) the shape of the displacement likelihood and parametrized it in the following way:

$$w_{fj0} = \begin{cases} \frac{1}{2}\left(\frac{d/dt - \mu_{fj}}{\sigma_{fj}}\right)^2 + c_{fj}\sqrt{dt} \, \frac{\arctan\left(\left|\frac{dy}{dx}\right|\right)}{\pi} & \text{if dx} < 0 \text{ and } |\text{dy}| < |\text{dx}| \\ 100 & \text{otherwise} \end{cases}$$

where $d$ is the displacement in time $dt$, $\mu_{fj} = 25\ um, \sigma_{sj} = 12.9um$ , and $c_{fj} = 36$.

$$w_{fj} = \frac{1}{\sqrt{2}}(w_{fj0} + c_{fj,t}(dt - 1)),$$

where $dt$ is segment duration in timeframes and $c_{sj,t} = 3$.

Since potential segments only with NLL below the not-connected NLL $w_{nc}$ can be selected by the optimization procedure, the segments with NLL above $w_{nc}$ were discarded. The remaining potential segments of the types “segmentation jump” and “flow jump” were appended to the graph (Figure 3E, Supplementary Figure 3B).

Next, we found the optimal multiplicity on each segment $M_{seg}$ and potential segment $M_{pseg}$. We identified T cell jumps along the track as the potential segments where the multiplicity $M_{pseg}$ was found to be non-zero. To this end we performed optimization under the constraint of global multiplicity consistency analogous to the one described above for the track segment search. Optimization was performed on each subgraph connected by segments independently. Here we demanded the same multiplicity on both ends of the segment $M_{seg,l} = M_{seg,r}$, consistent with corresponding multiplicity on the connected vertex $M_{seg[i],r} = M[i]_{vtx,l}$, and same multiplicity on both sides of the vertex,

$$M_{end\_l} + M_{nc_l} + \sum\nolimits_{i,l} M_{seg}[i] + \sum\nolimits_{i,l} M_{pseg}[i] = M_{end\_r} + M_{nc_r} + \sum\nolimits_{i,r} M_{seg}[i] + \sum\nolimits_{i,r} M_{pseg}[i],$$

where $M_{end}$ is the multiplicities in case the vertex is a track endpoint, $M_{nc}$ is the resolved missing multiplicity at the vertex, and $M_{seg}[i], M_{pseg}[i]$ are the multiplicities of the attached to the vertex segments and potential segments on left and right sides. Left and right endpoint multiplicity $M_{end\_l/r} = 0$ if any segment is connected on the corresponding side. We constrained $0 \le M_{nc} \le M_{max}$, $f_0 \le M_{seg}[i] \le M_0 + M_{max}$, and $0 \le M_{pseg}[i] \le M_{max}$ where $M_{max} = 5$. We did not consider intersections of more than 5 T cells, as those could not be reliably identified on the opposite ends of the intersections by their motility parameters (see the intersection resolving section), and in our experiments we found this value to be sufficient.

Optimization was subjected to minimizing the loss function in each node:

$$L_n = \sum_{l,r} M_{end} * w_{end} + \sum_{l,r} M_{nc} * w_{nc} + \sum\nolimits_{i} (M[i] - M_0[i]) * w_{over} + \sum\nolimits_{i} c_{ps}[i] * w_{ps}[i]$$

Here we distinguished missing multiplicity on a vertex $M_{nc} \neq 0$, corresponding to T cells detaching in a group of under-segmented T cells when some of the T cell tracks continued further from the endpoint of the track, where $M_{end} \neq 0$. The first term of the loss function penalized higher multiplicity at the endpoints of the track, encouraging solutions with T cell tracks being properly segmented in the endpoints. Here $M_{end}$ is the multiplicity at the corresponding endpoint, and the $w_{end} = 6$.

Putting additional weight on the missing multiplicity on a vertex in the middle of the graph but not on the first node of a track as opposed to all nodes at non-first timeframes of the dataset allowed us to account for flow causing T cell tracks to start and end at any timeframe due to T cell attachment during the accumulation phase and detachment of T cells carried away by the flow. Thus, the second term penalizes the missing multiplicity on either side of the vertex with the coefficient $w_{nc}$:

$$w_{nc} = f_{edge}\, f_{time} * w_{nc\_0},$$

where $w_{nc\_0} = 9$, $f_{edge}$ corresponds to the attenuation close to the edges of the FoV as described above. We considered the T-cell accumulation phase under low flow by attenuating the weight of track start and end $w_{nc}$ with time (Supplementary Figure 4E):

$$f_{time,l} = \begin{cases} \frac{1}{9} & \text{if } t < t_{acc.start} \\ \frac{1}{9} + \frac{8}{9}\frac{t - t_{acc.start}}{t_{acc.end} + 5 - t_{acc.start}} & \text{if } t_{acc.start} \le t < t_{acc.end} + 5 \\ 1 & \text{otherwise} \end{cases}$$

$$f_{time,r} = \begin{cases} \frac{1}{2} & \text{if } t < t_{acc.end} \\ \frac{2}{9} & \text{if } t_{acc.end} \le t < t_{acc.end} + 5 \\ 1 & \text{otherwise} \end{cases}$$

This corresponded to a higher probability for track to start or end (T cell attaching/detaching from the pMBMECs monolayer) during the accumulation phase, as well as a high probability for the track to end during the 5 frames after the flow was increased to the physiological level.

The third term of the loss function encouraged minimal adjustments to the multiplicity estimates by penalizing the difference between the initial multiplicity estimate on T cell track segments and the final solution with $w_{over} = 1$.

The last term accounted for the potential segments. $c_{ps}[i] = 1$, if the multiplicity $M_{pseg}$ on the corresponding potential segment was found to be above zero, and $c_{ps}[i] = 0$ when the multiplicity was zero. $w_{ps}[i]$ – is the NLL of the potential segment, i.e., either $w_{fj}$ or $w_{sj}$ for the flow- or segmentation-caused jump segments correspondingly. All the coefficients were chosen empirically to optimize the detection of T-cell jumps.

Finally, we eliminated short and thus unreliable segments, as well as segments for which the multiplicity was found to be $M = 0$ (Figure 3F). To this end, we iteratively removed vertices, which were not connected on one side and connected to a branching vertex of the graph on the other side by a segment with less than 6 nodes, as well as the connecting segment, and repeated the global multiplicity optimization until reaching convergence. To reduce the wall time required for the optimization, it was again performed independently for each connected component of the dataset graph.

Afterwards, we separated the segments into two categories, namely segments with multiplicity $M = 1$, i.e., tracks of isolated T cells, and segments with multiplicity $M > 1$ i.e. tracks of under-segmented groups of T cells, where tracks of several T cells intersected (Figure 3G).

In the end, we extended the tracks of isolated T cells into the intersection if the branching vertex had a multiplicity of $n$ and was splitting in exactly $n$ segments with multiplicity $M = 1$ (Figure 3G). Specifically, we performed the segment extension if there was an unambiguous assignment of cells constituting the branching vertex to the segments connected to the vertex, i.e., that the weight $w_{i,j}$ of assigning $i$-th cell belonging to the vertex to $j$-th segment connected to the vertex satisfies simultaneously $\arg\min_j w_{i_a,j} = j_a$ and $\arg\min_i w_{i,j_a} = i_a$. To this end, we evaluated the standard deviation $\sigma_j$ of the distance from the T cell along the $j$-th segment to the line obtained by linear extrapolation of the cell coordinated at the nearest 6 timeframes. We then obtained the expected T-cell position $\widehat{r_{j,t}}$ by linear extrapolation of the $j$-th segment to the timeframe $t$ of the vertex using the nearest 6 timeframes of the segment. The assignment weight was evaluated as the NLL:

$$w_{i,j} = \frac{1}{2}\left(\frac{dr_{i,j}}{\sigma_j}\right)^2,$$

where $dr_{i,j} = r_{i,t} - \widehat{r_{j,t}}$ is the distance between position $r_{i,t}$ of the $i$-th cell belonging to the vertex at timeframe $t$, and the expected cell position $\widehat{r_{j,t}}$.

*Intersection resolving*

The last step required to obtain reliable T cell tracks is resolving track intersections, i.e., identifying track segments corresponding to the same T cell before and after the under-segmented track region (Figure 3H). Even though T-cell motility parameters were found to be quite similar within the population investigated, evaluating them on longer segments allowed us to identify the correspondence between segments of the same T-cell track.

To this end, we evaluated the NLL of potential assignments of track segments according to the distance between their closest points and mean T cell crawling speed (cell track path length over time), migration speed (cell displacement over time), directionality, cell ellipticity, and cell area. We estimated these parameters employing the Bayesian estimator with prior expected values evaluated on tracks of isolated T cells obtained after resolving the global track consistency. We then used F-statistic to assess the connection probability and to obtain the corresponding NLL. We then employed the Linear Assignment Problem (LAP) approach to identify the optimal segment assignments across the track intersections. To ensure that reliable connections were made, instead of relying on the assignment with the first solution for all tracks, we instead connected only the segments with the $\mathrm{NLL} < \mathrm{NLL}_{max,i}$ progressively increasing $\mathrm{NLL}_{max,i}$ in $\{1, 3, 6, 9, \infty\}$ and repeating the parameter estimation for the remaining unresolved track segments and the LAP solving. This ensured that the T-cell motility parameters used for segment matching were estimated on the longer, already connected T-cell track segments.

This workflow allowed us to determine fully resolved T-cell tracks, with either assigned T-cell position or identified locations of rapid T-cell displacement, cell under-segmentation timespan, and missing time-frames due to potential T-cell detection inefficiency (Figure 3H).

#### ***3. T-cell migration analysis***

Next, we performed the T-cell migration analysis based on the reconstructed T-cell tracks. We selected tracks inside the fiducial area of the FoV, namely coordinates of the T cell at all timepoints along the track were located at least 25 µm away from the bounding box enclosing all segmented T cells. Next, tracks of T cells touching another T cell at the end of the assay acquisition were excluded since T cells directly adjacent to each other can hide the start of T cell transmigration across the pMBMEC monolayer and thus compromise correct detection and quantification of this step. Additionally, only tracks with T cells assigned in at least 6 timeframes during the physiological flow phase. We also require T cells to be assigned for at least 75% of timeframes along the track. Under-segmented parts of T cell tracks were not considered. Examples of selected tracks can be seen in Supplementary Video 3.

First, we counted the number of neighboring tracks, i.e., T cells approaching another T cell closer than 24 µm (1.2 of a crawling T cell length).

T-cell transmigration across the pMBMEC monolayer was detected based on the inferred T-cell transmigration coefficient $t_c$. To reduce the noise, we obtained filtered time-series of transmigration coefficients $t_{c,f}$ by applying a Gaussian filter with $\sigma = 2.5$ timeframes to the $t_c$ time-series. We defined the T cell to the preliminary categories "full transmigration" and category "partial transmigration" with $t_{c,f} > 0.75$ and $t_{c,f} > 0.3$ respectively. The "full transmigration" and "partial transmigration" Boolean masks were constructed according to the evolution of these two categories along the track (Supplementary Figure 5). To further reduce the noise, we also closed short gaps (sequence of False values) in the transmigration masks with length 2 and 3 timeframes for the "partial" and "full" transmigration respectively. Then we removed short sequences of True values in the masks of 4 and 3 timeframes correspondingly.

Based on these preliminary masks, we detected the additional categories of uncompleted transmigration, direct transmigration, and reverse transmigration categories (Figure 1D). During uncompleted transmigration, the T cell starts the transmigration process and later on retracts the protrusions to continue crawling above the pMBMEC monolayer. Direct transmigration covers the period during which the T cell transmigrates below the pMBMEC monolayer. In contrast, during reverse transmigration, the T cell previously located below the pMBMEC monolayer reversely migrates back to the luminal side of the pMBMEC monolayer (Supplementary Figure 5). The start, duration, and number of transmigration attempts were evaluated for each transmigration category.

We employed an additional classifier to discriminate a migrating T cell from cellular debris or particles (=not-a-T-cell) to reject the cellular debris misclassified as T cells during the segmentation step. This classification was performed based on the measured mean and median T cell areas along the T cell tracks after the flow was increased to physiological levels. Specifically, we trained a logistic regression model using the scikit-learn package (6). The model was trained on 105 manually annotated T cells by an experimenter and subsequently validated on 27 additional T cells. We have achieved validation accuracy of 96%. The tracks classified as "not-a-T-cell" were discarded from the downstream analysis.

Due to occasional over-segmentation of the T cell during the transmigration process, a T cell could be misclassified as detaching instead of as transmigrated. To address this, we have trained a linear classifier to differentiate a detached T cell from a transmigrated T cell. In this case, the classification was performed based on the following parameters: mean $a_{mean}$

and median $a_{med}$ T cell area along the part of the T-cell track during the physiological flow phase, $\frac{a_{last\ 1}}{a_{mean}}, \frac{a_{last\ 1}}{a_{med}}, \frac{a_{mean\ last\ 5}}{a_{mean}}, \frac{a_{mean\ last\ 5}}{a_{med}}$, where $a_{last\ 1}$ and $a_{mean\ last\ 5}$ were the T-cell area at the last and mean over the last 5 timeframes of the T cell track correspondingly, mean $p_{c,mean}$ and median $p_{c,med}$ T-cell detection probability as well as probabilities at the last timeframe $p_{c,last\ 1}$ and mean of last 5 timeframes $p_{c,mean\ last\ 5}$, and the mean transmigration coefficient over the last 5 timeframes of the T cell track $t_{c,mean\ last\ 5}$.

If the track of a T cell remaining above the pMBMEC monolayer ended before the end of the assay and this T cell is classified as detaching, it was marked as a detached T cell accordingly. Following their transmigration across the pMBMEC monolayer, T cells continued to crawl below the pMBMEC monolayer in the microfluidic device but were no longer subjected to flow. If such tracks were ending before the end of the assay, they were labeled as "tracking inefficiency" and were subsequently excluded from further analysis of the post-transmigration motility parameters.

Next, we created masks of tracking inefficiency and periods of accelerated movement – i.e., rapid displacement of the T cells on the pMBMEC monolayer caused by the flow. For each T-cell track during the physiological flow phase and prior to transmigration we evaluated the median step speed $v_{step,med}$, i.e., the T-cell speed between sequential timeframes, and its median absolute deviation $MAD(v_{step})$. We evaluated the mean instantaneous speed $v_{inst,mean}$ and its standard deviation $\sigma_{v,inst}$, excluding steps where the speed was an outlier, i.e., $v_{inst} > v_{inst,med} + 3\ \sigma_{v\ inst\ med}$, where $\sigma_{v\ inst\ med} = 1.4826\ MAD(v_{inst})$. We labeled timespan along the T cell tracks as "accelerated T cell movement" if the instantaneous speed

$$v_{inst} > \min\left(v_{inst,mean} + 3\ \sigma_{v,inst}, v_{crawling\ max}\right)$$

and the instantaneous displacement $dr_{inst} > dr_{am}$, where $v_{crawling\ max} = 36\ \mu m/min$ and $dr_{am} = 8\ \mu m$. If a step had a time difference of more than 3 timeframes, e.g., in the case of intersecting T cell tracks – it was labeled as tracking inefficiency.

If the T-cell displacement was more than 20 $\mu m$ (length of a crawling T cell) during a cell track segment, irrespective if above or below the pMBMEC monolayer, the segment was marked as crawling. This is analogous to the criteria used during manual analysis (see the "Materials and Methods" section). The cell track was marked as probing if the T cell did not crawl before the first transmigration attempts.

Finally, for each track, we evaluated motility parameters for each of the following migration regimes: probing before the transmigration, crawling before the transmigration, all crawling above pMBMECs monolayer including T cell crawling segments after the first transmigration attempts, all crawling below pMBMECs monolayer, whole T-cell track excluding accelerated movement and tracking inefficiency regions, as well as whole T-cell track. Specifically, we evaluated the following T-cell motility parameters: duration of each migration regime, the total vector and absolute displacements, the migration path length, the average migration speed (displacement over time), average crawling speed (path length over time), and finally the mean and standard deviation of the instantaneous speed. For the accelerated movement regime, we evaluated migration time, displacement, and average speed.

**Supplementary Table 1**: Architecture of the 2D fully convolutional model for T cell segmentation.

| # | Operation | Kernel | Stride | DO rate | Output size | Diagram |
|---|---|---|---|---|---|---|
| 1 | Input | | | | 256 × 256 × 1 | |
| 2 | Conv+LReLU | 7 × 7 | | | 256 × 256 × 64 | |
| 3 | Conv+BN+DO+LReLU | 3 × 3 | | 0.2 | 256 × 256 × 64 | |
| 4 | MP | | 2 × 2 | | 128 × 128 × 64 | |
| 5 | Conv+BN+DO+LReLU | 3 × 3 | | 0.2 | 128 × 128 × 128 | |
| 6 | Conv+BN+DO+LReLU | 3 × 3 | | 0.2 | 128 × 128 × 128 | |
| 7 | MP | | 2 × 2 | | 64 × 64 × 128 | |
| 8 | Conv+BN+DO+LReLU | 3 × 3 | | 0.2 | 64 × 64 × 256 | |
| 9 | Conv+BN+DO+LReLU | 3 × 3 | | 0.2 | 64 × 64 × 256 | |
| 10 | MP | | 2 × 2 | | 32 × 32× 256 | |
| 11 | Conv+BN+DO+LReLU | 3 × 3 | | 0.2 | 32 × 32× 512 | |
| 12 | TConv+BN+LReLU | 3 × 3 | 2 × 2 | | 64 × 64 × 256 | |
| 13 | Concat(12, 9) | | | | 64 × 64 × 512 | |
| 14 | Conv+BN+DO+LReLU | 3 × 3 | | 0.2 | 64 × 64 × 256 | |
| 15 | Conv+BN+DO+LReLU | 3 × 3 | | 0.2 | 64 × 64 × 256 | |
| 16 | TConv+BN+LReLU | 3 × 3 | 2 × 2 | | 128 × 128 × 128 | |
| 17 | Concat(16, 6) | | | | 128 × 128 × 256 | |
| 18 | Conv+BN+DO+LReLU | 3 × 3 | | 0.2 | 128 × 128 × 128 | |
| 19 | Conv+BN+DO+LReLU | 3 × 3 | | 0.2 | 128 × 128 × 128 | |
| 20 | TConv+BN+LReLU | 3 × 3 | 2 × 2 | | 256 × 256 × 64 | |
| 21 | Concat(20, 3) | | | | 256 × 256 × 128 | |
| 22 | Conv+BN+DO+LReLU | 3 × 3 | | 0.2 | 256 × 256 × 64 | |
| 23 | Conv+BN+DO+LReLU | 3 × 3 | | 0.2 | 256 × 256 × 64 | |
| 24 | └ Conv+Sigmoid | 1 × 1 | | | 256 × 256 × 2 | |
| 25 | └ Conv+BN+DO+LRe | 3 × 3 | | 0.1 | 256 × 256 × 16 | |
| 26 | Conv+SSigmoid | 3 × 3 | | | 256 × 256 × 1 | |
| 27 | Concat(24, 26) | | | | 256 × 256 × 3 | |

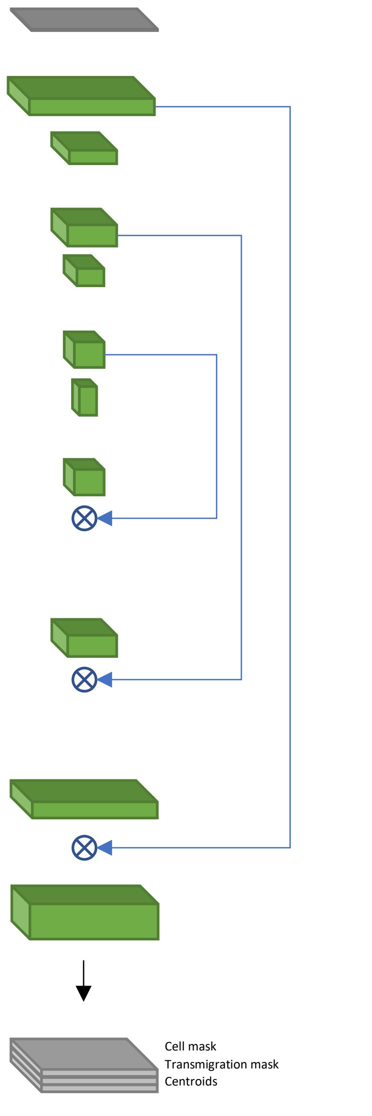


DO – dropout, Conv – Convolution, TConv – transposed convolution, MP – max pooling, BN – batch normalization, Concat – concatenation of outputs of the specified layers, LReLU – leaky rectified linear unit with α=0.05.

For the centroids prediction, we scale the output to prevent saturation: SSigmoid(x) = 1.1 * Sigmoid(x) – 0.5.

**Supplementary Table 2**: Architecture of the 2D+T fully convolutional model for T cell segmentation

| # | Operation | Kernel | Stride | DO rate | Output size | Diagram |
|---|---|---|---|---|---|---|
| 1 | Input | | | | 5 × 256 × 256 × 1 | |
| 2 | Conv+LReLU | 1 × 7 × 7 | | | 5 × 256 × 256 × 64 | |
| 3 | Conv+BN+DO+LReLU | 3 × 3 × 3 | | 0.2 | 5 × 256 × 256 × 128 | |
| 4 | └ ZCrop | | | | 1 × 256 × 256 × 128 | |
| 5 | └ ZCrop | | | | 3 × 256 × 256 × 128 | |
| 6 | MP | | 1 × 2 × 2 | | 3 × 128 × 128 × 128 | |
| 7 | Conv+BN+DO+LReLU | 1 × 3 × 3 | | 0.2 | 3 × 128 × 128 × 128 | |
| 8 | Conv+BN+DO+LReLU | 1 × 3 × 3 | | 0.2 | 3 × 128 × 128 × 128 | |
| 9 | └ ZCrop | | | | 1 × 128 × 128 × 128 | |
| 10 | └ MP | | 1 × 2 × 2 | | 3 × 64 × 64 × 128 | |
| 11 | Conv+BN+DO+LReLU | 1 × 3 × 3 | | 0.2 | 3 × 64 × 64 × 256 | |
| 12 | Conv+BN+DO+LReLU | 3 × 3 × 3 | | 0.2 | 3 × 64 × 64 × 512 | |
| 13 | ZCrop | | | | 1 × 64 × 64 × 512 | |
| 14 | MP | | 1 × 2 × 2 | | 1 × 32 × 32 × 512 | |
| 15 | Conv+BN+DO+LReLU | 1 × 3 × 3 | | 0.2 | 1 × 32 × 32 × 256 | |
| 16 | Conv+BN+DO+LReLU | 1 × 3 × 3 | | 0.2 | 1 × 32 × 32 × 256 | |
| 17 | MP | | 1 × 2 × 2 | | 1 × 16 × 16 × 256 | |
| 18 | Conv+BN+DO+LReLU | 1 × 3 × 3 | | 0.2 | 1 × 16 × 16 × 512 | |
| 19 | TConv+BN+LReLU | 1 × 3 × 3 | 1 × 2 × 2 | | 1 × 32 × 32 × 256 | |
| 20 | Concat(19, 16) | | | | 1 × 32 × 32 × 512 | |
| 21 | Conv+BN+DO+LReLU | 1 × 3 × 3 | | 0.2 | 1 × 32 × 32 × 256 | |
| 22 | TConv+BN+LReLU | 1 × 3 × 3 | 1 × 2 × 2 | | 1 × 64 × 64 × 128 | |
| 23 | Concat(22, 13) | | | | 1 × 64 × 64 × 256 | |
| 24 | Conv+BN+DO+LReLU | 1 × 3 × 3 | | 0.2 | 1 × 64 × 64 × 128 | |
| 25 | Conv+BN+DO+LReLU | 1 × 3 × 3 | | 0.2 | 1 × 64 × 64 × 128 | |
| 26 | TConv+BN+LReLU | 1 × 3 × 3 | 1 × 2 × 2 | | 1 × 128 × 128 × 128 | |
| 27 | Concat(26, 9) | | | | 1 × 128 × 128 × 256 | |
| 28 | Conv+BN+DO+LReLU | 1 × 3 × 3 | | 0.2 | 1 × 128 × 128 × 128 | |
| 29 | Conv+BN+DO+LReLU | 1 × 3 × 3 | | 0.2 | 1 × 128 × 128 × 128 | |
| 30 | TConv+BN+LReLU | 1 × 3 × 3 | 1 × 2 × 2 | | 1 × 256 × 256 × 64 | |
| 31 | Concat(30, 4) | | | | 1 × 256 × 256 × 192 | |
| 32 | Conv+BN+DO+LReLU | 1 × 3 × 3 | | 0.2 | 1 × 256 × 256 × 64 | |
| 33 | Conv+BN+DO+LReLU | 1 × 3 × 3 | | 0.2 | 1 × 256 × 256 × 64 | |
| 34 | └ Conv+Sigmoid | 1 × 1 × 1 | | | 1 × 256 × 256 × 2 | |
| 35 | └ Conv+BN+DO+LReLU | 1 × 3 × 3 | | 0.1 | 1 × 256 × 256 × 16 | |
| 36 | Conv+SSigmoid | 1 × 3 × 3 | | | 1 × 256 × 256 × 1 | |
| 37 | Concat(34, 36) | | | | 1 × 256 × 256 × 3 | |

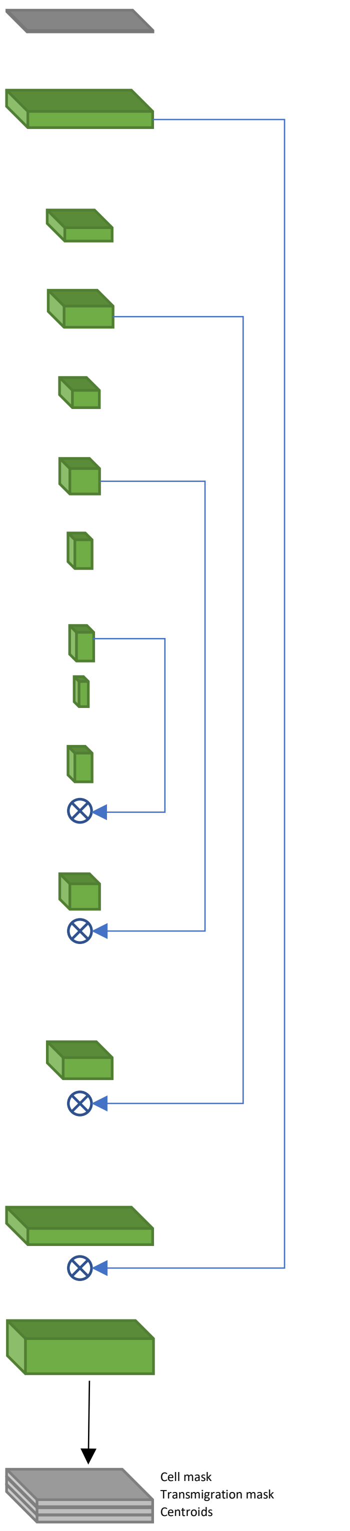


DO – dropout, Conv – Convolution, TConv – transposed convolution, MP – max pooling, BN – batch normalization, Concat – concatenation of outputs of the specified layers, ZCrop – crop center along depth dimension, LReLU – leaky rectified linear

unit with $\alpha = 0.05$.

For the centroids prediction, we scale the output to prevent saturation: SSigmoid(x) = 1.1 * Sigmoid(x) – 0.5.

**Supplementary Table 3:** Node variables used in global optimization during link search.

| Name | Description | Type | Range |
|---|---|---|---|
| $c_{l/r}[i]$ | Multiplicity of the [i]-th node connection on the left/right side | Int | 0..m |
| $c_{w\_l/r}[i]$ | i-th connection on the left/right is selected | Int | 0..1 |
| $c_{nc\_l/r}$ | Not connected (NC) connection on the left/right is selected | Int | 0..1 |
| $d_{abs\_lr}$ | Absolute difference between right and left multiplicity sum | Int | 0.. m |

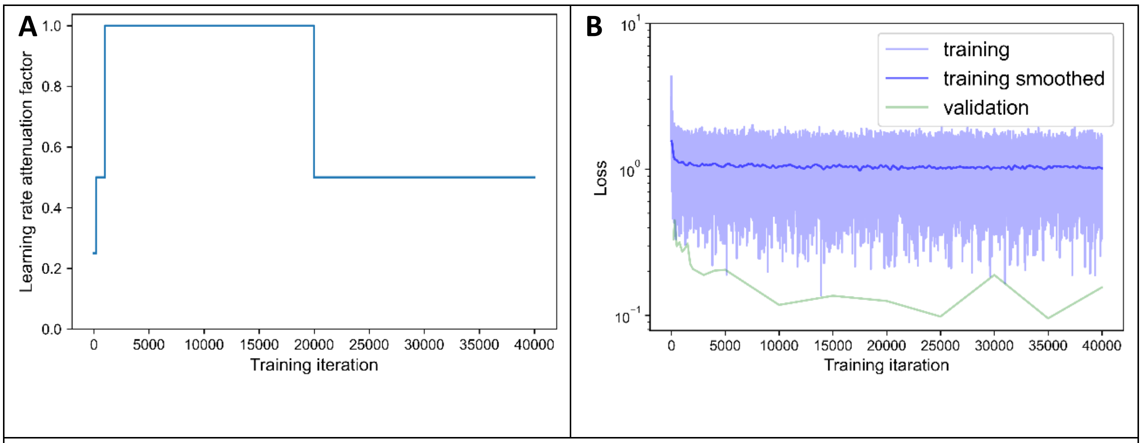


**Supplementary Figure 1. Training curves**. **A.** Learning rate attenuation along model training. **B.** Loss value evolution along model training.

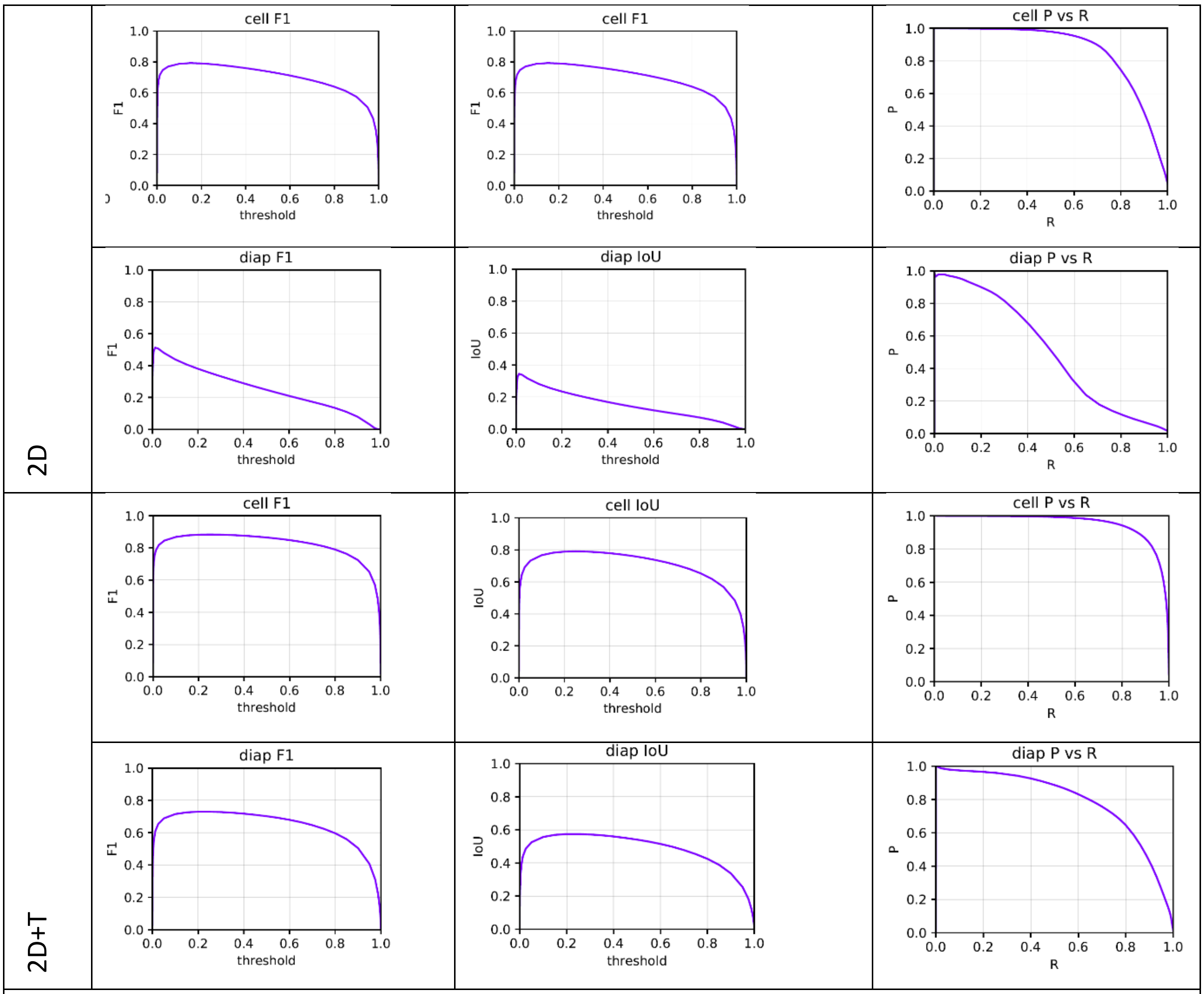


**Supplementary Figure 2. Comparison of the 2D and 2D+T cell segmentation models' performance for the T cell mask and transmigration mask prediction.** F1, Jaccard index, and AP metrics are shown.

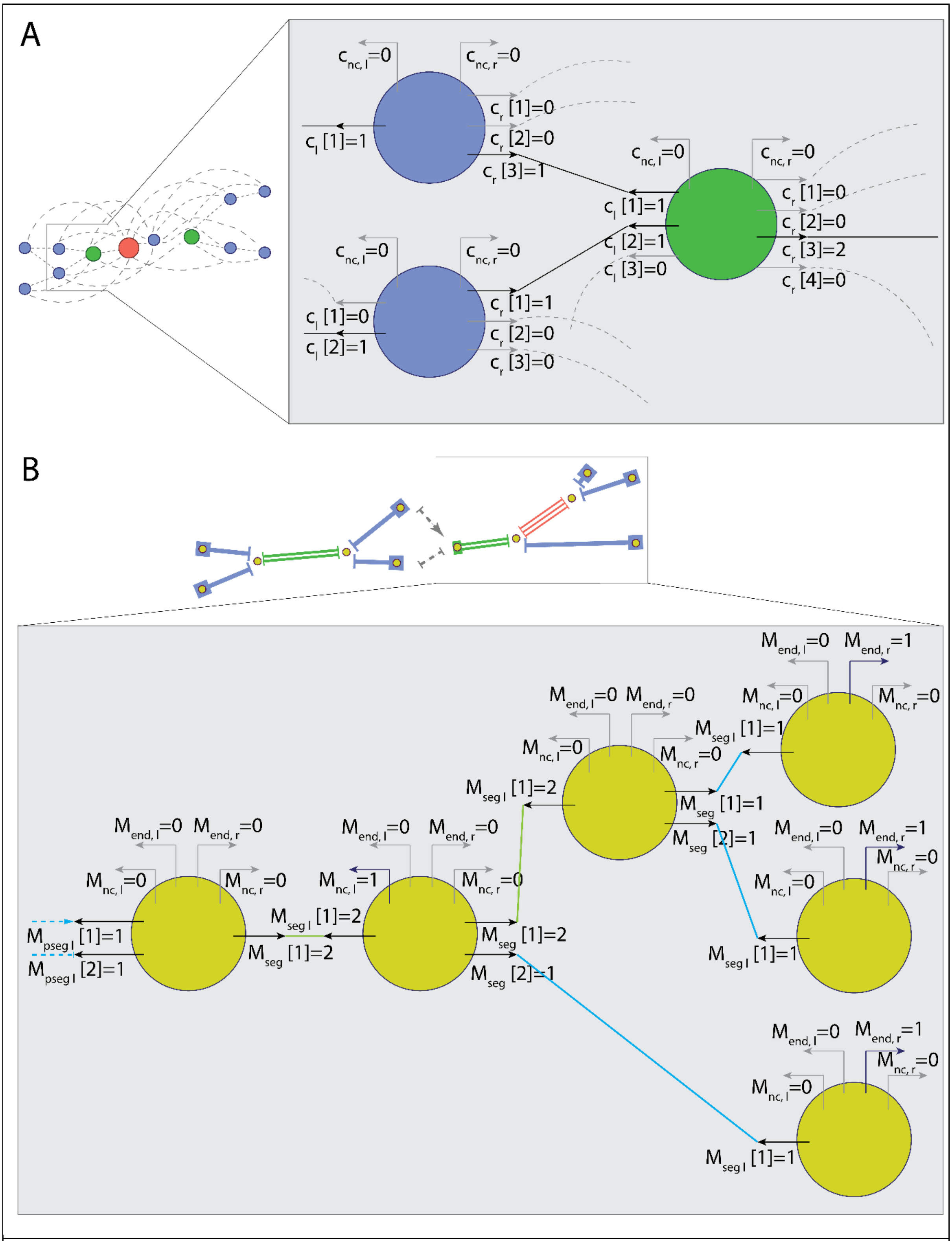


**Supplementary Figure 3: Parametrization of nodes connections for tracking. A.** Representation of nodes for track segment search. Connection multiplicity for each link is obtained with global optimization. **B.** Representation of vertices and segments for global multiplicity consistency optimization. Connection multiplicity for each segment attached to a vertex is obtained with global optimization.

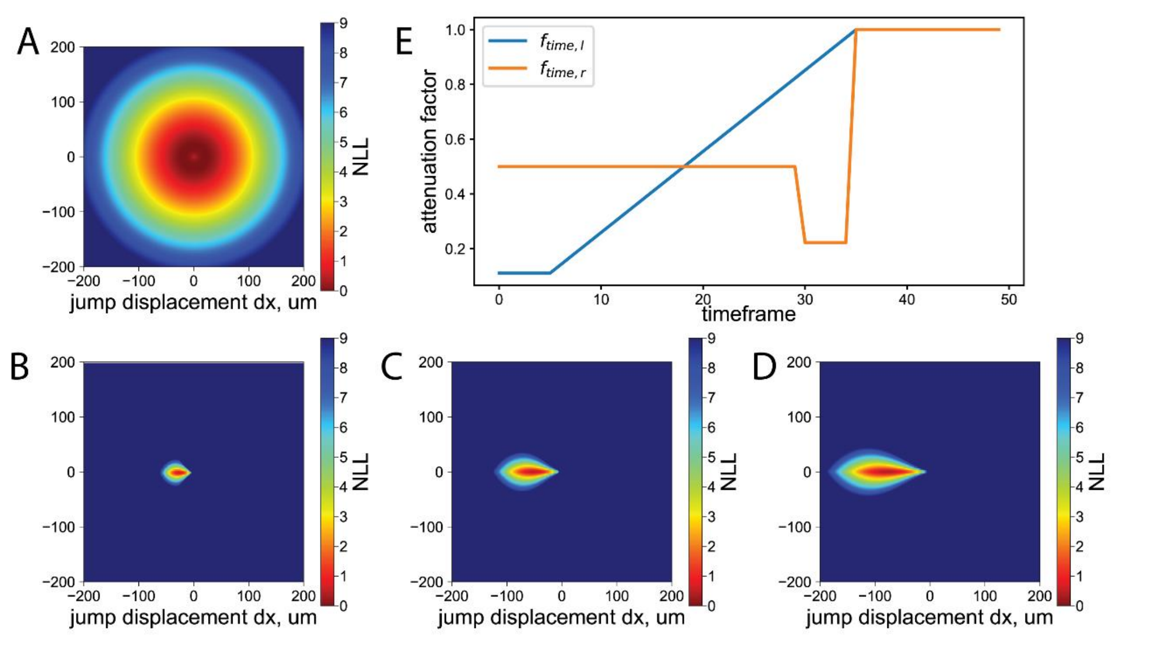


**Supplementary Figure 4: Underflow tracking parametrization. A**. Negative log-likelihood $w_{sj0}$ of the potential T cell jump segments due to under-segmentation. **B-D.** Negative log-likelihood $w_{fj0}$ of the potential T cell jumps segments due to the flow. **B**: dt=1, **C**: dt=2, **D**: dt=3. **E.** Attenuation of the vertex "not-connected" weight with time allows accounting for the T-cell accumulation phase at timeframes 5 through 30 and increase of the flow to physiological level at timeframe 30. The blue curve shows an attenuation factor on the left side, i.e., corresponding to track start, and the orange curve shows an attenuation factor on the right side corresponding to the end of the track due to cell detachment under the flow.

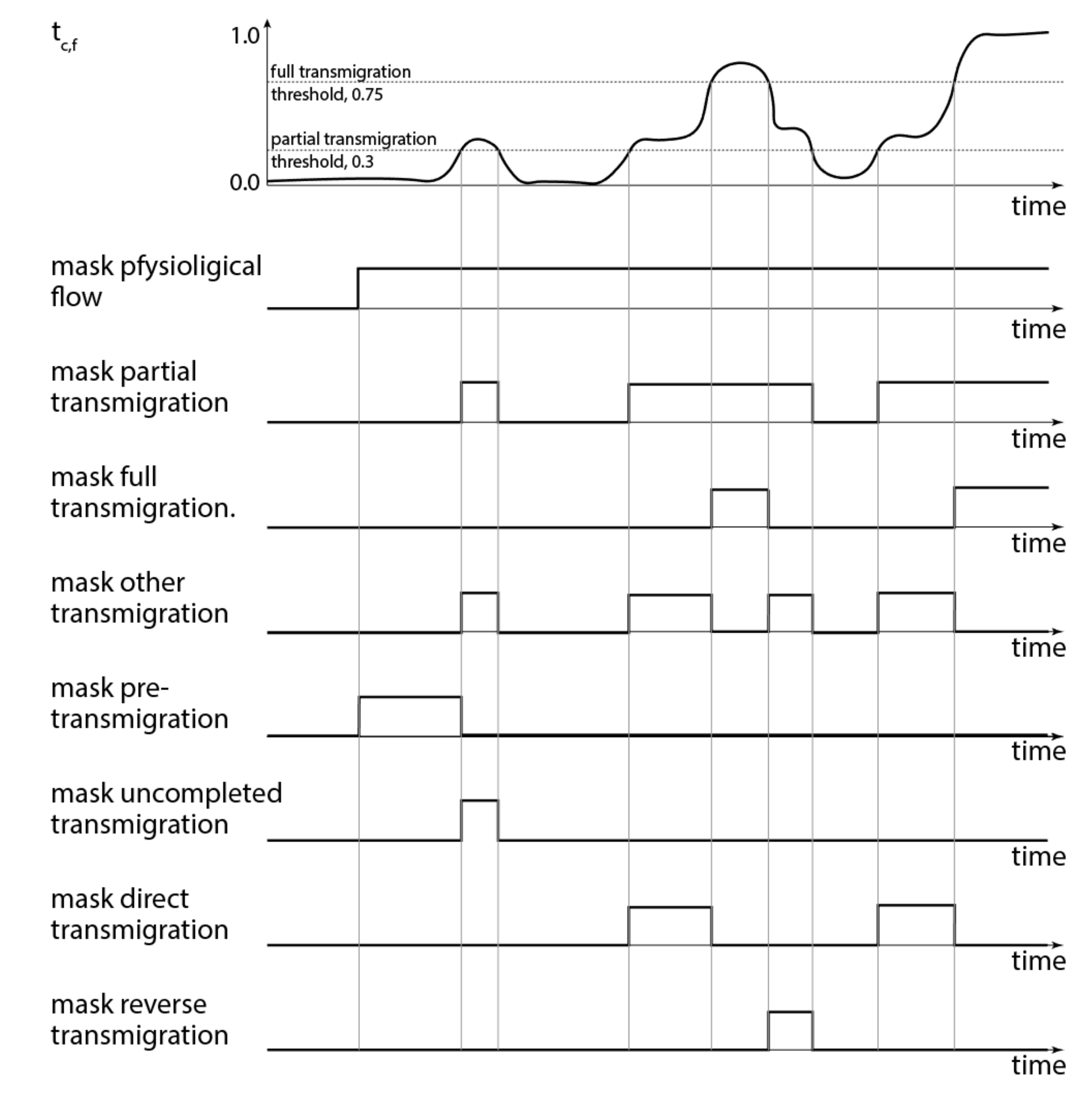


**Supplementary Figure 5. Transmigration detection.** Based on the filtered transmigration factor $t_{c,f}$ we obtained Boolean masks for partial and full transmigration. Next, we obtained Boolean masks for T cell migration before and during uncompleted, direct, full, and reverse transmigrations.

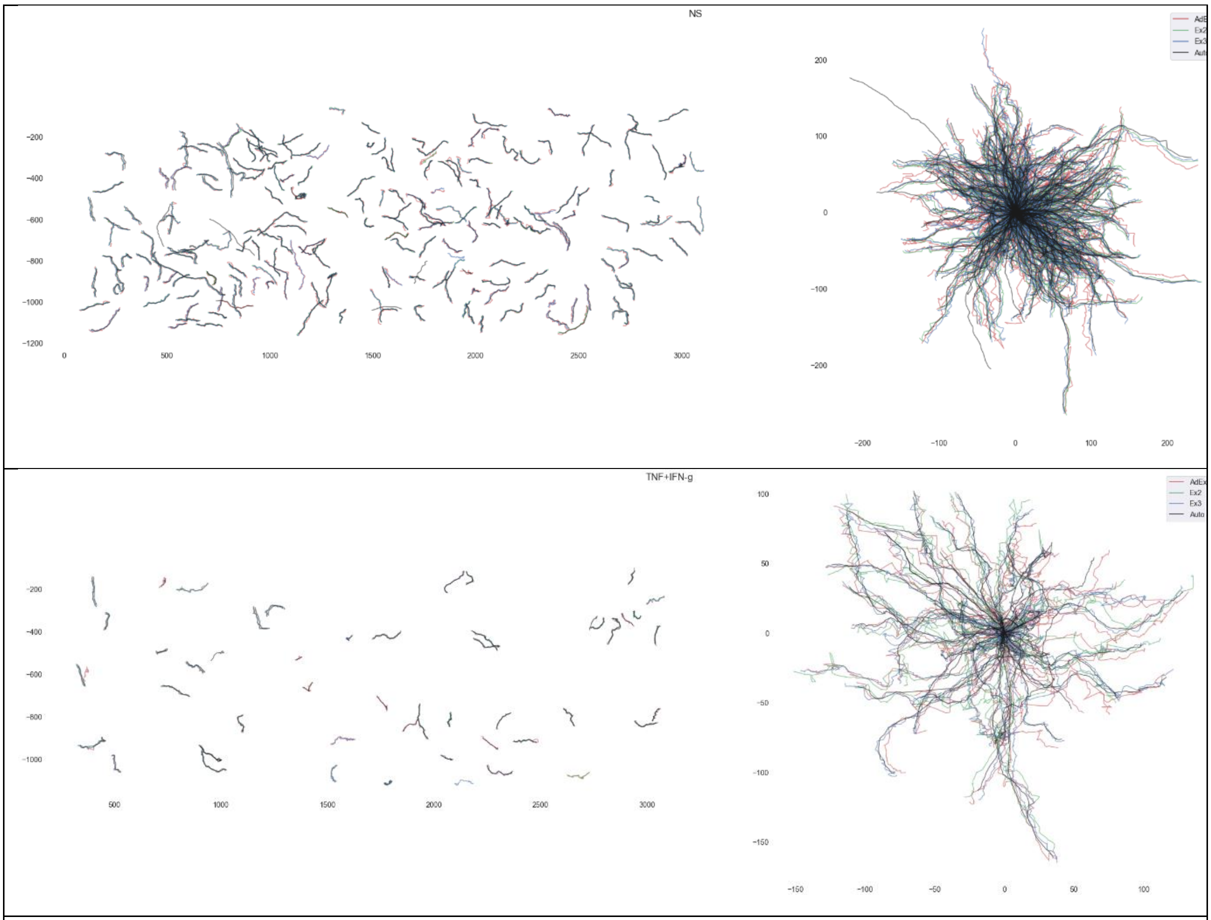


**Supplementary Figure 6. Comparison of reconstructed T cell tracks with the result of manual analysis.** Overlay of crawling CD8$^{+}$ T-cell crawling tracks on top of non-stimulated (top) and TNF/IFNγ stimulated pMBMECs (bottom) tracked manually by three experimenters as well as automatically. Errors in manual tracking led to a jittery pattern in the T-cell tracks. This lead, in turn to the overestimation of the T cell crawling speed.

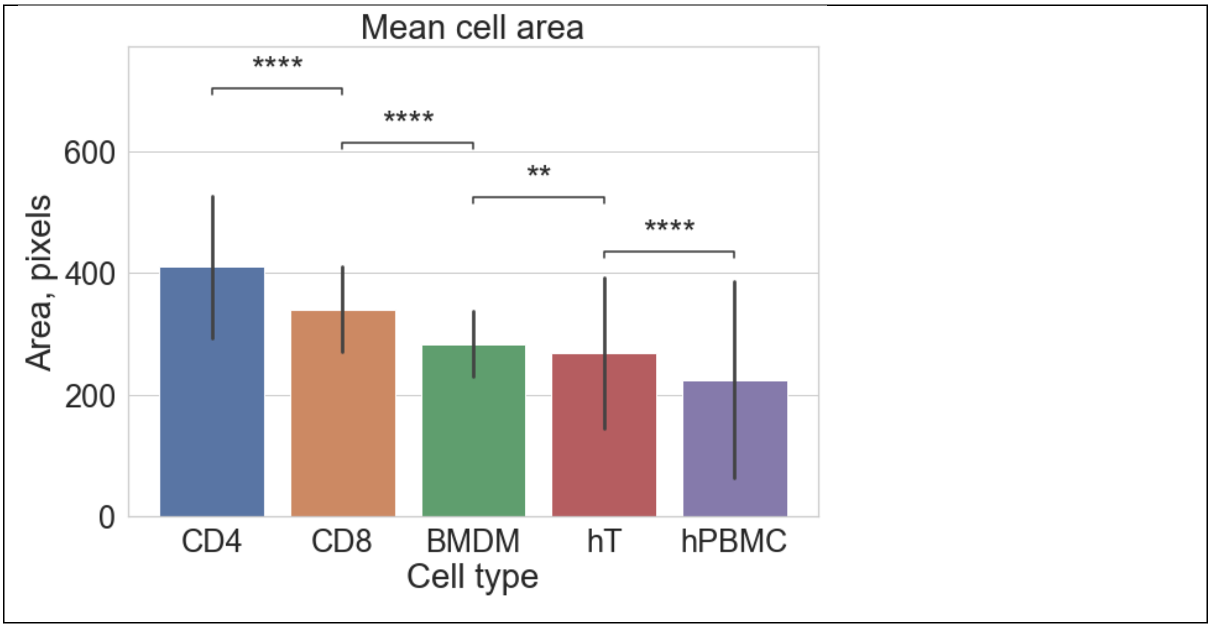

**Supplementary Figure 7. Comparison of sizes of crawling cells.** $^{UFM}$Track is able to successfully reconstruct tracks of migrating cells of different sizes. CD4 and CD8 – mouse $CD4^+$ and $CD8^+$ cells, BMDM – bone marrow derived macrophages, hT – human T cells, hPBMC – human peripheral blood mononuclear cells.

**Supplementary Video 1.** Phase-contrast time-lapse image sequence of CD4 T cells interacting with IL-1β stimulated endothelium. 8 tiles of the imaging are aligned and stitched together.

**Supplementary Video 2.** Segmented T cells in the phase-contrast time-lapse image sequence of CD4 T cells interacting with IL-1β stimulated endothelium. The mask of the segmented T cell is overlayed in red. The transmigration probability map is overlayed in yellow.

**Supplementary Video 3.** Tracks of T cells reconstructed in the phase-contrast time-lapse image sequence of CD4 T cells interacting with IL-1β stimulated endothelium. Tracks are shown after the flow increase to a shear stress level of 1.5 dynes/cm2. All videos are shown accelerated by a factor of 96 as can be seen on the timestamp label. Only tracks included in the analysis are shown (see text for details).
